# Sequential evolution of antidote and toxin links genetic incompatibility with immune responses

**DOI:** 10.64898/2026.01.07.698274

**Authors:** Dongying Xie, Yiming Ma, Junhui Zeng, Pohao Ye, Zhongying Zhao

## Abstract

Toxin-antidote (TA) systems are selfish genetic elements that promote their own inheritance by selectively eliminating offspring lacking the module, thereby establishing post-zygotic genetic incompatibilities between individuals and populations. Although these systems are widespread across species, their evolutionary origins remain poorly understood. Here, we report the discovery of a novel TA gene pair in the nematode *Caenorhabditis nigoni*. The antidote gene, *Cni-shls-2*, is a species-specific F-box gene that arose through recent tandem duplications. Its absence results in embryonic lethality in both *C. nigoni* and its hybrids with the sister species *C. briggsae*. This lethality is caused by a maternally deposited toxin, *Cni-hlix-1*, a chimeric gene formed through the fusion of host and bacterial sequences. Phylogenetic and genomic analyses reveal a sequential evolution of the TA system, in which the antidote evolved prior to the toxin. The stepwise evolution of TA and the potential microbial origin of the toxin support the hypothesis that the antidote initially evolved in response to pathogen exposure, followed by the domestication of the toxin, thereby elucidating the origins of TA formation. These findings highlight the central role of host-pathogen conflict as a driving force in the emergence of genetic incompatibilities and the evolution of reproductive barriers.

## Introduction

Organisms face constant intrinsic and extrinsic challenges that drive genomic divergence and the evolution of new genetic modules, ultimately contributing to the formation of genetic incompatibilities among individuals. Selfish genetic elements, particularly a specific type known as toxin-antidote (TA) systems, exemplify one such incompatibility (*1*). Comprising a toxic agent and its corresponding detoxifying component, TA systems bias inheritance by impairing gametes or offspring that lack both elements and consequently generate reproductive barriers (*2*). While TA systems are pervasive and well-characterized in prokaryotes (*3*), their occurrence and impact have only recently been reported in eukaryotes, including fungi (*4–7*), nematodes (*8–13*) and plants (*14, 15*). This has spurred speculation about their potentially more universal existence than previously recognized, even including vertebrates (*1*). Nevertheless, a fundamental question remains largely unresolved: how do these selfish genetic elements originate in the first place?

The inception of TA systems is central to understanding their prevalence and mechanisms (*1, 13*). Given that the toxin inherently impairs offspring fitness, it seems unlikely that it would evolve independently. Conversely, the evolution of an antidote gene, which may confer no immediate adaptive benefits on its own, raises questions about its initial utility. Two models have been proposed to explain the origins of TA systems (*1*). The endogenous model suggests that TA systems emerge as intrinsic adaptive mechanisms, supported by the discovery that multiple nematode toxin genes exhibit homology with some conserved essential genes (*9, 13*). A recent study even suggests that the recurrent formation of TA pairs may originate from pre-suppressive modules that buffer against deleterious effects of random mutations in essential genes (*13*). In contrast, the exogenous model posits that TA systems may arise from host-pathogen arms races, as suggested by TA elements derived from intracellular bacterial parasites, for example, *Wolbachia*, which trigger cellular incompatibilities in hosts such as *Drosophila* (*16, 17*). However, the mechanism by which endogenous TA elements arise in the host genome remains poorly understood.

The abundance of hybrid incompatibilities between the sister nematode species pair, *Caenorhabditis briggsae* and *C. nigoni*, provides an ideal opportunity to dissect both the evolution of genetic conflict and the origins of TA systems (*18–20*). Our previous work demonstrates an asymmetrical gene flow between the two species (Fig. 1A), largely attributable to the presence of multiple *C. nigoni-*specific *shls* (<u>s</u>pecies <u>h</u>ybrid lethality <u>s</u>uppressor) genes, whose loss frequently leads to detrimental hybrids (Fig. 1B) (*20, 21*). Given the potential prevalence of TA elements in *Caenorhabditis* nematodes (*11*), many of which lead to embryonic lethality, we reason that at least a subset of these *shls* genes may function as antidotes within selfish TA modules. Here, we report the identification of a novel TA element in *C. nigoni* that underlies embryonic lethality upon its absence both within the species and in hybrid offspring with *C. briggsae*. These findings suggest that the antidote likely originated in response to selective pressures from the host immune system against external pathogens, followed by the domestication of the toxin. Our results underscore the significant role of host-pathogen arms races in driving the emergence of genetic incompatibilities and the establishment of reproductive barriers.

**Figure 1.**
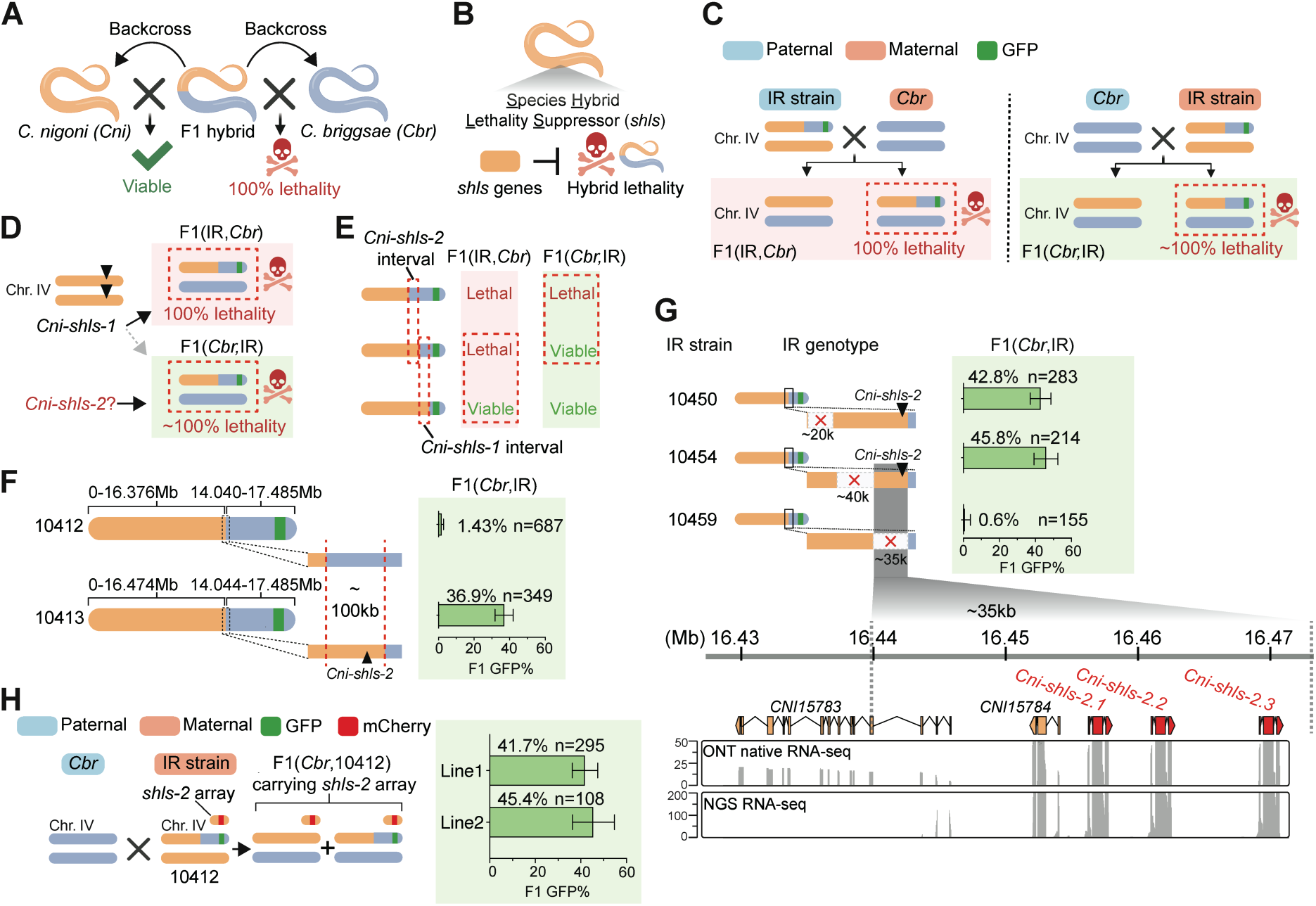
A newly identified suppressor gene underlines hybrid lethality in crosses between a *C. briggsae* father and a *C. nigoni* mother. **(A)** Schematic illustrating asymmetrical lethality for backcross hybrids between *C. briggsae* and *C. nigoni*. Hybrid F1 progeny of the two species yield viable and fertile offspring when backcrossed to *C. nigoni*, but no viable progeny when backcrossed to *C. briggsae*. **(B)** The indispensability of some *C. nigoni* genomic regions indicates that *C. nigoni* harbors <u>s</u>pecies <u>h</u>ybrid lethality <u>s</u>uppressor (*shls*) gene(s) essential for the viability of hybrids of the two species. **(C)** Schematic showing a parent-of-origin independent hybrid lethality phenotype observed when a *C. nigoni* introgression (IR) strain carrying a heterozygous *C. briggsae* Chr. IV right arm fragment is crossed with *C. briggsae*. F1 hybrids are named with the paternal strain on the left of the comma and the maternal strain on the right within parentheses. **(D)** Schematic illustrating that the recently identified gene *Cni-shls-1* suppresses lethality in hybrids fathered but not mothered by *C. briggsae*, implying a potential second suppressor gene, *Cni-shls-2*. **(E)** Schematic showing that in F1 hybrids, homozygosity for larger or smaller *C. briggsae* Chr. IV right arm introgression results in lethality or viability, respectively, whereas an intermediate introgression causes lethality only when *C. briggsae* is the mother, confirming separate target intervals encompassing *Cni-shls-1* and *Cni-shls-2*. **(F) Left:** Schematic comparing the genomic differences between two IR strains, ZZY10412 and ZZY10413. **Right:** Bar plot comparing the percentage of GFP-expressing adult F1 hybrids when the two IR strains are crossed with a *C. briggsae* father. Error bars represent 95% confidence intervals. **(G) Top:** Systematic deletion of the ∼100kb interval in the ZZY10413 background and the corresponding percentage of GFP-expressing F1 adults when crossed with *C. briggsae* fathers. **Bottom:** Predicted gene models within the ∼35 kb candidate interval, with RNA-seq read coverage (ONT and NGS) from *C. nigoni* embryos shown below. **(H) Left:** Schematic of a cross between a *C. briggsae* father and a ZZY10412 mother carrying the *Cni-shls-2* gene as an extrachromosomal array. **Right:** Bar plot showing the percentage of GFP-expressing adult F1 hybrids that carry the *Cni-shls-2* array. Two independent array lines were tested. Error bars represent 95% confidence intervals.

## Results

### Genetic mapping and cloning of a second hybrid lethality suppressor gene in *C. nigoni*

To systematically identify *shls* genes in *C. nigoni*, we previously generated a large cohort of *C. nigoni* strains, each carrying a heterozygous, GFP-linked introgression of a *C. briggsae* genomic fragment (hereafter referred to as IR strains) (*19, 20*). We reasoned that, in crosses between each IR strain and *C. briggsae*, half of the F1 hybrid progeny would be homozygous for the *C. briggsae* introgression, thereby lacking the corresponding syntenic *C. nigoni* region. If the missing region contains a *shls* gene, the affected hybrids would be inviable. Consistent with this, we identified several candidate regions (*20*), among which the right arm of chromosome IV (Chr. IV) was particularly notable: its absence resulted in complete hybrid lethality independent of the parent-of-origin (Fig. 1C) (*20, 21*). We recently cloned the first underlying *shls* gene, *Cni-shls-1* (*21*). Intriguingly, however, *Cni-shls-1* is required only for the viability of hybrids from *C. briggsae* mothers, as maternally deposited *Cni-shls-1* is sufficient to sustain the development of hybrids lacking zygotically expressed SHLS-1 protein (Fig. 1D) (*21*). Consistently, during our systematic mapping of *Cni-shls-1* using IR strains with variable introgression lengths, we observed a shift from parent-of-origin-independent to - dependent lethality (Fig. 1E and Table S1) (*21*). Specifically, homozygosity for relatively larger or smaller *C. briggsae* introgression fragments resulted in hybrid lethality or viability, respectively, as evidenced by the absence or presence of GFP-expressing hybrid F1 adults, whereas an intermediate introgression caused lethality only in hybrids derived from *C. briggsae* mothers (Fig. 1E, Fig. S1 and Table S1). These findings support the existence of an additional *shls* gene, i.e., *Cni-shls-2*, whose absence leads to lethality in hybrid F1 progeny from *C. briggsae* fathers.

To further narrow the candidate interval for *Cni-shls-2*, we focused on two key IR strains, ZZY10412 and ZZY10413, both of which were generated using our recently developed targeted recombination method (*21, 22*) (Fig. 1F). Although both strains exhibited hybrid lethality when crossed with *C. briggsae* mothers (Fig. S1), ZZY10412 displayed hybrid lethality when crossed with *C. briggsae* fathers, as evidenced by the almost absence of GFP-expressing hybrid adults (1.43%), whereas ZZY10413 did not (36.9%) (Fig. 1F). ONT (Oxford Nanopore Technology) long-read sequencing revealed a critical difference in their recombination boundaries: ZZY10413 (0-16.474Mb) retains approximately 100 kb more of the *C. nigoni* genomic fragment compared to ZZY10412 (0-16.376Mb) (Fig. 1F and Fig. S2). In contrast, their corresponding *C. briggsae* genome breakpoints differ by only about 4 kb (14.040-17.219Mb in ZZY10412 vs. 14.044-17.219Mb in ZZY10413) (Fig. 1F and Fig. S2). We therefore reasoned that the absence of the ∼100 kb *C. nigoni* region in ZZY10412, where *Cni-shls-2* resides, rather than the minor difference in the *C. briggsae* fragment, alters hybrid viability.

The ∼100 kb candidate region contains 14 predicted genes (Fig. S3). To further narrow down the interval, we subsequently performed three large genomic fragment deletions for the chromosome with the *C. briggsae* introgression in ZZY10413 via CRISPR/Cas9, such that each deletion interval encompassed 4-5 predicted genes (Fig. S3). We then tested the viability (presence/absence of GFP-expressing adults) of the hybrids between the deletion mutant mother and *C. briggsae* father (Fig. 1G). Deletion of either the left ∼20 kb (43.8%) or the middle ∼40 kb region (45.8%) yielded a slightly higher percentage of GFP-expressing adult hybrids than that from ZZY10413 (36.9%) (Fig. 1G). However, the removal of the right ∼35 kb region resulted in a remarkable drop in the proportion of GFP-expressing F1 adults (0.6%) (Fig. 1G). This result confirms that *Cni-shls-2* resides in the rightmost segment of the ∼100kb region, which contains four predicted genes, i.e., *CNI15784*, *CNI15785*, *CNI15786*, and *CNI15787* (Fig. 1G and Fig. S4A). Notably, the four genes are separated by abundant tandem repeats of the same type (Fig. S5A), and *CNI15785*, *CNI15786*, and *CNI15787* are tandem duplicates with identical coding sequences, albeit with slight variations in their introns and upstream regions (Fig. S5A and B).

To pinpoint *Cni-shls-2*, we generated additional deletions either targeting *CNI15784* or the genomic region encompassing *CNI15785* to *CNI15787*, on the *C. nigoni* Chr. IV carrying the *C. briggsae* introgressed fragment in ZZY10413 (Fig. S4B-D). We then assessed hybrid viability of these mutant strains when crossed with *C. briggsae* fathers. Hybrid inviability was observed only when the region containing the three tandemly duplicated genes (*CNI15785*-*CNI15787*) was deleted (Fig. S4E-G), demonstrating that these genes encode the *Cni-shls-2* locus. To confirm this, we performed a complementation test by introducing an extrachromosomal array containing a single copy of the genomic sequences of *Cni-shls-2* into strain ZZY10412 (Fig. 1H). In contrast to nearly complete hybrid lethality observed in the absence of the array, the presence of the array from two independent lines restored GFP-expressing hybrid viability to 41.7% and 45.4%, supporting that zygotic expression of *Cni-shls-2* is sufficient for the rescue of hybrid lethality (Fig. 1H). We therefore molecularly cloned the second *shls* gene and named the three identical copies as *Cni-shls-2.1*, *Cni-shls-2.2*, and *Cni-shls-2.3*.

### *Cni-shls-2* is a *C. nigoni*-specific F-box gene in the same family as *Cni-neib-1*

To investigate the function of *Cni-shls-2*, we first generated a mutant allele by deleting all three paralogous copies in the wild-type *C. nigoni* background (Fig. 2A). We found that *Cni-shls-2* is essential for *C. nigoni* viability as homozygous mutations could not be obtained in viable animals and those carrying the homozygous mutations appeared to arrest during embryogenesis (Fig. 2A). Importantly, introducing an extrachromosomal array carrying *Cni-shls-2* into homozygous mutants rescued the lethality (Fig. 2B), indicating that zygotic expression of *Cni-shls-2* is essential for the viability.

**Figure 2.**
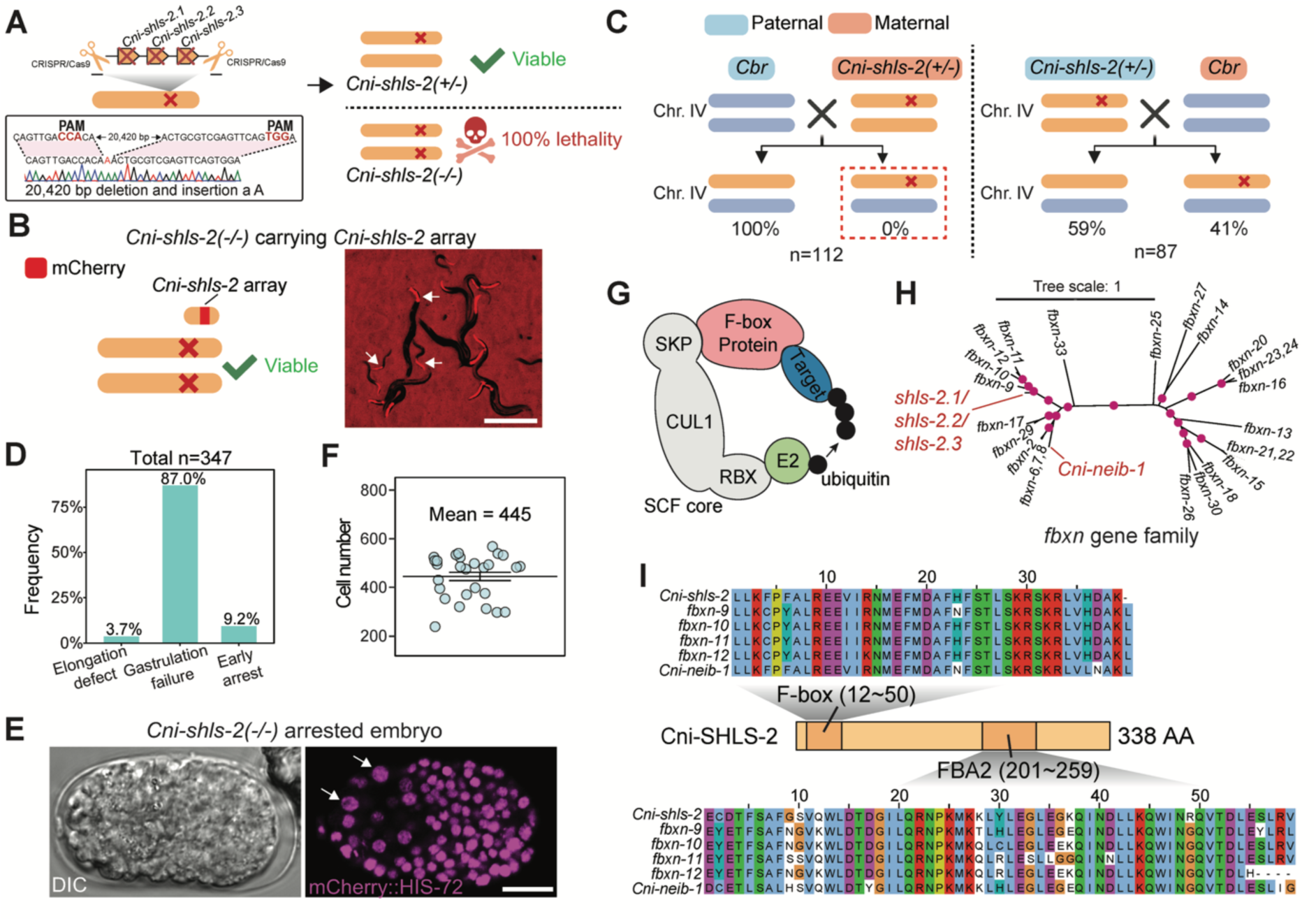
*Cni-shls-2* encodes an F-box protein within the same family of Cni-NEIB-1. **(A) Left:** Schematic representation and deletion junction of a *Cni-shls-2* mutant allele generated by CRISPR/Cas9. **Right:** Schematic demonstrating the inviability of *Cni-shls-2(-/-)* mutants. **(B) Left:** Schematic illustrating that a *Cni-shls-2* extrachromosomal array can rescue the inviability of *Cni-shls-2(-/-)* mutants. **Right:** Representative image of rescued homozygous mutants showing *pmyo-2::mCherry* expression (array marker) in the pharynx (arrowheads). Scale bar: 0.5 mm. **(C)** Comparison of the percentage of hybrid progeny carrying the *Cni-shls-2* deletion allele when crossing a *C. briggsae* father with a *Cni-shls-2(+/-)* mother versus the reciprocal cross. Note that *Cni-shls-2* is indispensable for hybrid viability only when the mutant allele is maternally inherited. **(D)** Bar plot showing the phenotype distribution of arrested *Cni-shls-2(-/-)* lethal embryos. **(E)** Representative image of a *Cni-shls-2(-/-)* lethal embryo that fails gastrulation. Aberrantly localized cells with enlarged nuclei are highlighted (white arrowheads). **(F)** Dot plot showing cell numbers in arrested *Cni-shls-2(-/-)* embryos. Error bars: SEM. **(G)** Schematic showing that F-box proteins act as adaptors by interacting with both the SCF E3 ubiquitin ligase complex and target substrates. **(H)** Protein phylogenetic tree of *fbxn* genes. Purple dots denote branches with bootstrap values >80. *Cni-neib-1* and *Cni-shls-2* are indicated. **(I)** Multiple sequence alignments of the F-box (top) and FBA-2 (bottom) domains from *Cni-shls-2*, its closely related *fbxn* genes (*fbxn-9* to *fbxn-12*), and *Cni-neib-1*.

We next assessed whether *Cni-shls-2* functions as a hybrid suppressor independent of the *C. briggsae* introgression fragment in a crossing direction-dependent manner. Reciprocal crosses between *Cni-shls-2* heterozygous mutants and *C. briggsae* revealed that no adult hybrids carried the deletion allele when a heterozygous *Cni-shls-2* mother was crossed with a *C. briggsae* father (0 out of 112) (Fig. 2C and Fig. S6). In contrast, nearly half of the progeny from the reciprocal cross carried the mutant allele (36 out of 87), consistent with Mendelian segregation (Fig. 2C and Fig. S6). These results confirm that the lack of *Cni-shls-2* is specifically responsible for the hybrid lethality between *C. briggsae* fathers and *C. nigoni* mothers in a parent-of-origin-dependent manner.

To understand how loss of *Cni-shls-2* affects embryogenesis, we examined arrested *Cni-shls-2(-/-)* embryos. Only a small fraction of embryos arrested at very early stages (with fewer than 28 cells) (9.2%) or at late stages when body elongation was evident (3.7%) (Fig. 2D). The majority (87%) exhibited an abnormal cell mass without obvious morphological organization, indicative of failed gastrulation (Fig. 2E). Consistent with this defect, we frequently observed large nuclei, presumably mislocalized gut cells at the periphery of embryos (Fig. 2E), although cell divisions continued with the majority of arrested embryos containing more than 300 cells (Fig. 2F). Collectively, these findings suggest that *Cni-shls-2* is crucial for proper gastrulation. To understand why *Cni-shls-2* is essential biochemically, we analyzed its encoded protein sequences. Surprisingly, rather than encoding a conserved gene like *Cni-shls-1*, *Cni-shls-2* is a *C. nigoni*-specific gene encoding an F-box protein. F-box proteins act as adaptors that link <u>S</u>kp1-<u>C</u>ullin-<u>F</u>-box (SCF) E3 ubiquitin ligase cores to substrate proteins for degradation (Fig. 2G). We previously identified a newly evolved, dispensable F-box gene, *Cni-neib-1*, in *C. nigoni* that fortuitously and specifically targets the highly conserved *C. briggsae Cbr-shls-1* gene, thereby causing embryonic lethality in hybrids lacking *Cni-shls-1* (*21*). Surprisingly, *Cni-shls-2* belongs to the same F-box gene family as *Cni-neib-1*, i.e., the *fbxn* (<u>F</u>-<u>b</u>o<u>x</u> *Cni-<u>n</u>eib-1*) gene family, which comprises 33 paralogous members mostly arranged in tandem (Fig. 2H), although each paralog displays varying degrees of protein sequence divergence (*21*). Besides *Cni-neib-1*, even the closest relatives of *Cni-shls-2*, i.e., *fbxn-9* to *fbxn-12* (Fig. 2H), exhibit considerable sequence dissimilarity (*21*). In addition, several amino acids differ in their F-box-associated domains (FBA2), the putative substrate recognition domain (*23*), supporting a differential function between *Cni-shls-2* and other *fbxn* members (Fig. 2I). Although F-box proteins can be co-opted for conserved function in nematodes (*24, 25*), it seems unlikely that *Cni-shls-2*, as a *C. nigoni*-specific F-box gene, performs similar roles essential for embryogenesis.

### *Cni-shls-2* encodes an antidote gene that mitigates the toxin activity of its cognate toxin gene *Cni-hlix-1*

We next asked why *Cni-shls-2*, a non-conserved, species-specific F-box gene, is indispensable for the development of both *C. nigoni* and its hybrids with *C. briggsae*. In nematodes, several species-specific genes have recently been identified as components of TA selfish elements (*9, 10, 12, 13, 26*). In these systems, one parent deposits toxin components into all zygotes, and individuals that do not inherit the TA allele lack the zygotically expressed antidote and are subsequently poisoned to death. Furthermore, recent studies have uncovered antidote-encoding genes as F-box genes in both nematodes (*13*) and rice (*15*). Given that *Cni-shls-2* is zygotically required for hybrid progeny when crossed with *C. briggsae* fathers, we hypothesized that *Cni-shls-2* encodes an antidote gene while a maternally deposited toxin from *C. nigoni*, which we named *Cni-hlix-1* for <u>h</u>ybrid lethality inducing to<u>x</u>in, kills progeny that do not inherit the TA gene pair (Fig. 3A).

**Figure 3.**
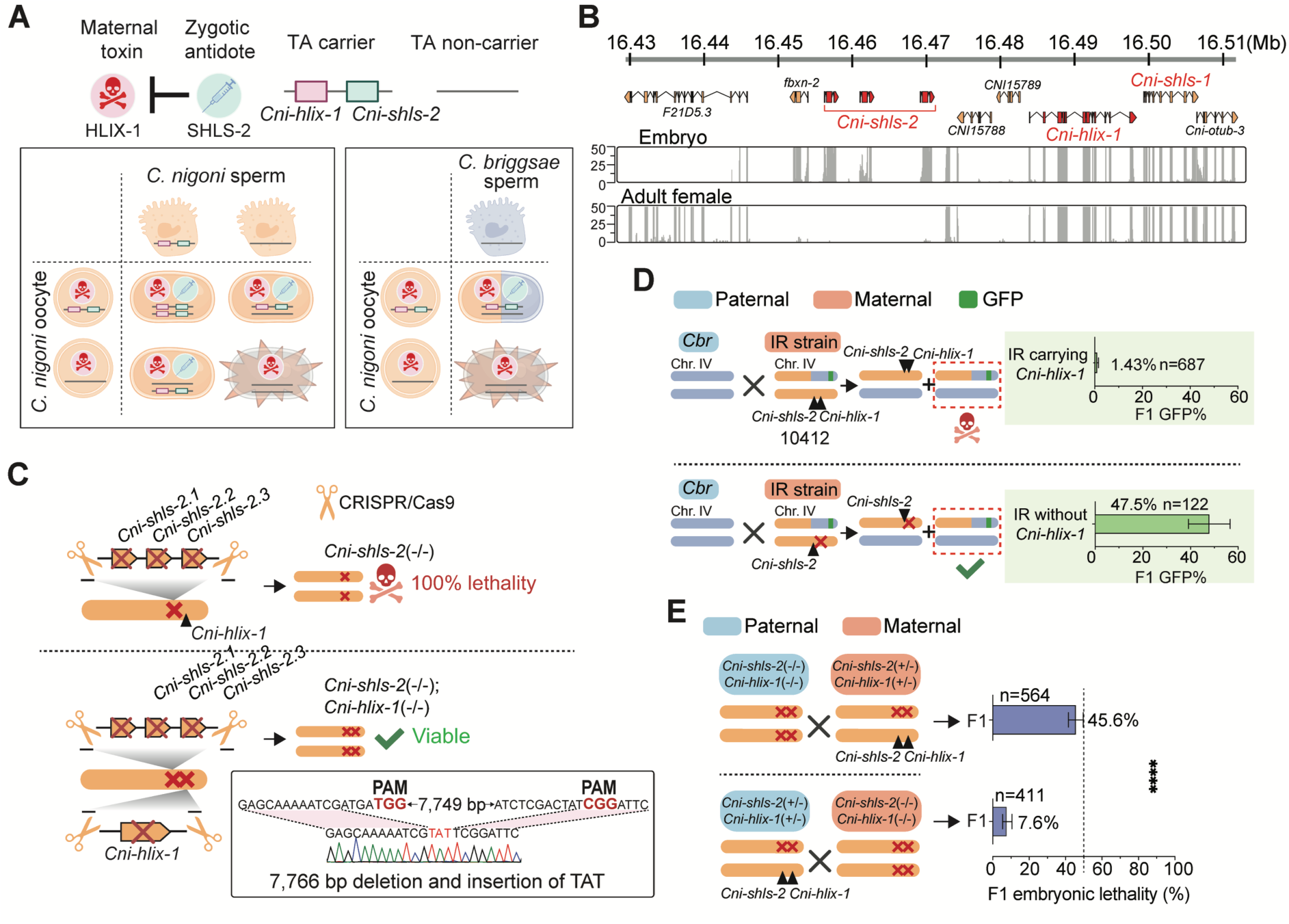
A novel TA gene pair underlies the embryonic lethality in *Cni-shls-2* mutants and their hybrids with *C. briggsae*. **(A)** Schematic illustrating the TA model, in which *Cni-shls-2* encodes the antidote and *Cni-hlix-1* encodes the toxin. Maternal deposition of Cni-HLIX-1 ensures that all zygotes inherit the toxin. Only those (*C. nigoni* or its hybrid with *C. briggsae*) that also inherit the TA element can neutralize its effect. **(B)** Predicted gene models adjacent to *Cni-shls-2*, with *Cni-hlix-1* and *Cni-shls-1* highlighted in red. RNA-seq read coverage (NGS) from *C. nigoni* embryos and adult females is shown below. **(C)** Schematic showing that *Cni-shls-2(-/-)* mutants are lethal (top), whereas *Cni-shls-2(-/-); Cni-hlix-1(-/-)* double mutants are viable (bottom). The CRISPR/Cas9-mediated deletion boundary of *Cni-hlix-1* is also shown. **(D)** Comparison of the percentage of GFP-expressing hybrid adults when crossing *C. briggsae* fathers with either a heterozygous IR strain mother carrying *Cni-hlix-1* (ZZY10412, top) or one carrying *Cni-hlix-1* mutant allele (bottom). Error bars: 95% confidence interval. **(E)** Box plot comparing embryonic lethality in F1 progeny from reciprocal crosses between homozygous and heterozygous double mutants of the TA gene pair. \*\*\*\**P*<0.0001, Fisher’s exact test. Error bars: 95% confidence interval.

To test this hypothesis, we first attempted to identify a potential toxin gene that is tightly linked to *Cni-shls-2* and is expected to be species-specific. We scrutinized the genomic region surrounding *Cni-shls-2.* After excluding three conserved genes (*F21D5.3*, *shls-1* (*pgm-3)*, and *otub-3*) as well as another *fbxn* gene, *fbxn-2*, three candidate genes were left: *CNI15788*, *CNI15789*, and *CNI15790* (Fig. 3B). RNA-seq analysis revealed that *CNI15790* is abundantly transcribed at both adult and embryonic stages, whereas the other two candidates displayed minimal expression at both stages (Fig. 3B). Thus, *CNI15790* is more likely to be *Cni-hlix-1* than the other two. To test this, we generated a *Cni-hlix-1* mutant in a *Cni-shls-2* homozygous mutant background (Fig. 3C), assuming that a double homozygous mutant of the putative TA element should be viable in the absence of toxin activity. As expected, the double homozygous mutant is viable and can be readily obtained (Fig. 3C). Next, to determine whether the hybrid lethality observed in progeny lacking *Cni-shls-2* is attributable to the toxicity of *Cni-hlix-1*, we first generated a *Cni-hlix-1* single mutant in the *C. nigoni* background that can also be maintained as a homozygote (Fig. S7A). We then crossed the *Cni-hlix-1* mutant with the IR strain ZZY10412 to ensure that a *Cni-hlix-1* mutant allele is present on the non-introgressed Chr. IV, followed by a cross with a *C. briggsae* father to assess hybrid lethality. In contrast to the ∼1.43% GFP-expressing hybrid progeny observed from a cross between ZZY10412 mothers and *C. briggsae* fathers, a significantly elevated proportion of GFP-expressing hybrids (47.5%) was observed from the cross between the IR mothers carrying a *Cni-hlix-1* null mutation and a *C. briggsae* father (Fig. 3D).

To further confirm that *Cni-hlix-1* is the toxin gene and the TA system functions via a maternal effect, reciprocal crosses were performed between double homozygous and double heterozygous mutants of *Cni-hlix-1* and *Cni-shls-2* (Fig. 3E). Consistent with a maternal toxic effect, when the heterozygous double mutant served as the mother (*Cni-hlix-1* was expected to be expressed and deposited maternally), approximately half of the progeny (45.6%) that did not inherit the TA genes (lacking the antidote) arrested during embryogenesis, whereas baseline levels of embryonic lethality (7.6%) were observed when the homozygous double mutant was the mother (no maternally deposited toxin) (Fig. 3E). A recent study demonstrated a beneficial effect of worms that carry another TA element (*peel-1/zeel-1*) in *C. elegans* (*27*). To assess whether the presence/absence of the TA element influences viability of *C. nigoni* or its hybrids with *C. briggsae*, we compared the hatching rates of wild-type *C. nigoni*, *Cni-hlix-1* homozygous mutants, and *Cni-hlix-1/Cni-shls-2* double homozygous mutants, as well as their hybrids with *C. briggsae* from reciprocal crosses (Fig. S7B and C). No significant differences in hatching rates were observed among wild-type *C. nigoni*, the single and the double mutants, suggesting that the presence of the TA pair minimally affects the fitness of *C. nigoni* and its hybrids with *C. briggsae*.

Collectively, we identified a novel pair of TA genes in *C. nigoni* that are both species-specific and tightly linked, with *Cni-hlix-1* encoding a maternally deposited toxin and *Cni-shls-2* encoding a zygotic antidote. The indispensability of *Cni-shls-2* for the embryonic viability of *C. nigoni* as well as its hybrid progeny with *C. briggsae* is therefore due to its detoxification effects.

### *Cni-hlix-1* is a chimeric gene formed by multiple gene reshuffling events

To understand how *Cni-hlix-1* kills embryos, we first examined its gene structure. ONT native RNA-seq reads confirmed the predicted model, revealing that *Cni-hlix-1* comprises 12 exons (Fig. S8). Although this gene is specific to *C. nigoni*, different segments share varying degrees of homology with other *C. nigoni* genes, indicating a chimeric origin (Fig. 4A). For example, exon 1 and a portion of exon 6 align in an inverse orientation to exon 2 of *Cni-prag-2*, a gene also located on Chr. IV, approximately 120 kb upstream of *Cni-hlix-1* (Fig. 4A and Fig. S9A and B). Moreover, the two genes upstream of *Cni-prag-2* (*CNI15773* and *CNI15774*) are highly homologous to those upstream of *Cni-hlix-1* (*CNI15788* and *CNI15789*) (Fig. S9A and B), and the sequences of *Cni-hlix-1* introns 1, 2 and 5 were largely derived from partial intron 1 of *Cni-prag-2*, which largely consists of simple tandem repeats (Fig. S9A and B). These results indicate that this part of *Cni-hlix-1* along with its upstream sequences arises from a segmental duplication followed by local rearrangement of *Cni-prag-2* genomic regions. The exons 1 and 6 of *Cni-hlix-1* appear to be split after their duplication from exon 2 of *Cni-prag-*2 (Fig. 4A and Fig. S9), which was caused by an insertion resulting from duplication and rearrangement of *Cni-lea-1* exons 4-7. They became exons 2-5 of *Cni-hlix-1* (Fig. 4A). *Cni-lea-1* is located on Chr. V, whose exonic structure appears to have arisen by tandem duplications (Fig. 4A and Fig. S9C). Finally, exons 8-12 and most of their corresponding intronic regions of *Cni-hlix-1* were derived from its downstream conserved gene, *Cni-otub-3*, an <u>o</u>varian <u>tu</u>mor (OTU) family deubiquitinase gene, as judged by the sequence homology (Fig. 4A and Fig. S9D). Aside from the C-terminal OTU domain, most Cni-HLIX-1 sequences are predicted to be intrinsically disordered regions (IDRs) (Fig. 4B).

**Figure 4.**
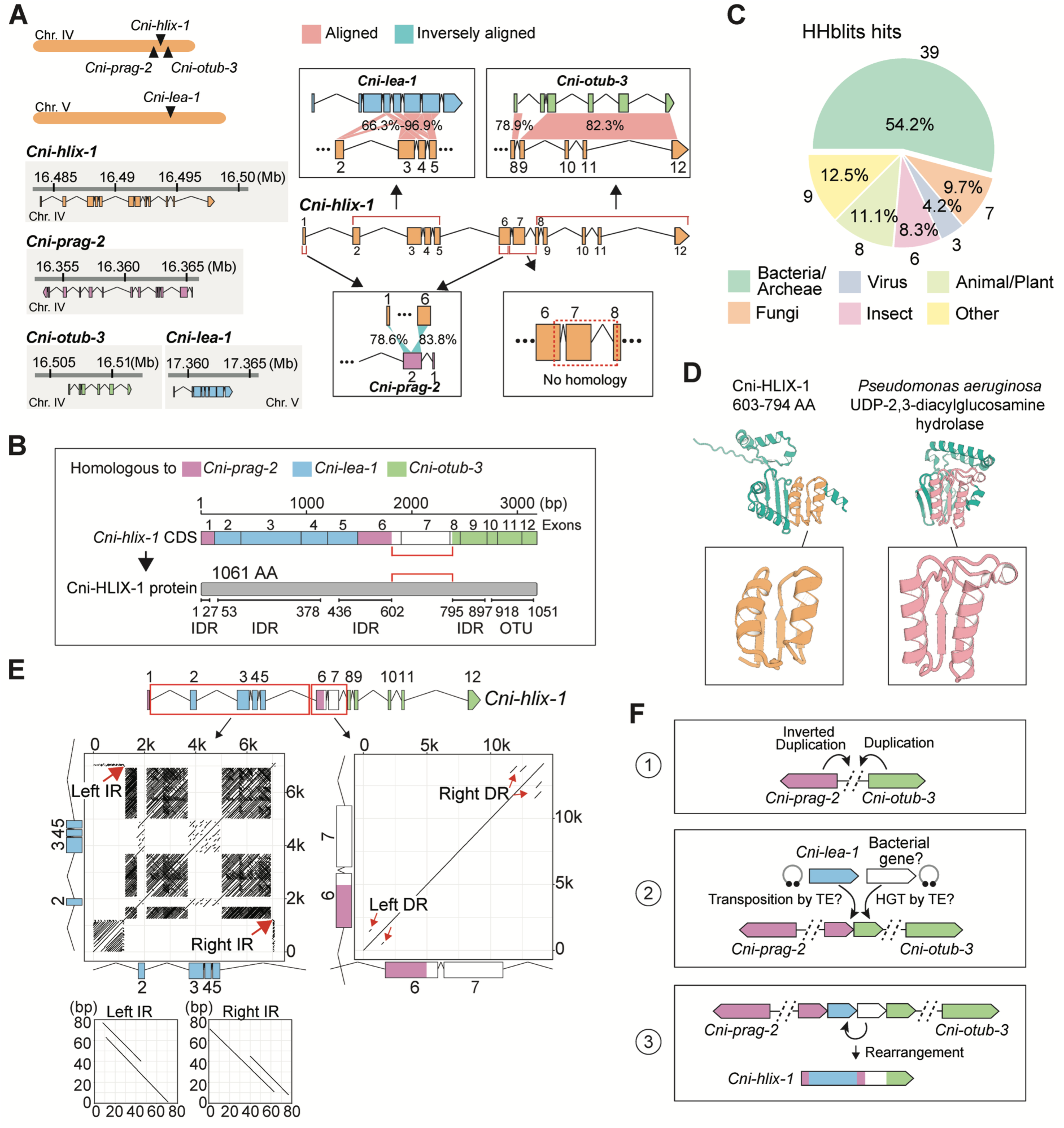
*Cni-hlix-1* is a chimeric gene formed through the fusion of host and bacterial sequences. **(A) Left:** Genomic positions and gene models of *Cni-hlix-1* and the genes that share homology to its segments: *Cni-lea-1* (blue), *Cni-prag-2* (purple), and *Cni-otub-3* (green). **Right:** Schematics showing the sequence homology between different parts of *Cni-hlix-1* and the three genes. Sequence similarity values are indicated. Portions of *Cni-hlix-1* without clear homology to nematode sequences are also indicated. **(B) Top:** Schematic of the portion of *Cni-hlix-1* CDS homologous to *Cni-lea-1*, *Cni-prag-2*, and *Cni-otub-3*. **Bottom:** Predicted protein domains of Cni-HLIX-1, with a region lacking clear homology and predictable domains (red bracket). IDR: intrinsically <u>d</u>isordered <u>r</u>egion; OTU: <u>o</u>varian <u>tu</u>mor deubiquitinase domain. **(C)** Pie chart showing the proportion of HHblits hits on different organisms for the no-homology region of Cni-HLIX-1 as highlighted in A and B. **(D)** Protein structure of the no-homology region in Cni-HLIX-1 (left) and that of the UDP-2,3-diacylglucosamine hydrolase from *Pseudomonas aeruginosa* (right). Structurally similar regions are highlighted. **(E)** Dot plots of the genomic regions between exons 1-6 (left) and exons 5-8 (right) of *Cni-hlix-1*. IR: inverted repeats; DR: direct repeats. Enlarged views of the left and right IRs in the region between exons 1-6 are shown below. **(F)** Schematic of the presumable evolutionary history of *Cni-hlix-1*.

Notably, one segment, comprising exon 7 and partial sequences from exons 6 and 8, shows no homology to any *C. nigoni* or nematode genes and lacks known protein domains (Fig. 4A and B). However, remote homology search returns over 50% bacterial or archaeal hits (Fig. 4C), albeit with relatively short alignments (Fig. S10), suggesting a potential bacterial origin. To further test this possibility, we used AlphaFold3 to predict the tertiary structure of this “no-homology” region and queried the Protein Data Bank (PDB). Although the N-terminal portion yielded few significant hits, the C-terminal region (∼50 amino acids) was highly similar in structure to segments of several bacterial enzymes such as hydrolases and helicases (Table S2). For instance, the 47-amino acid segment of the 192-aa no-homology region can be aligned to the N-terminal region of UDP-2,3-diacylglucosamine hydrolase from *Pseudomonas aeruginosa* (Fig. 4D). These observations demonstrate that *Cni-hlix-1* may have arisen from a fusion of both endogenous *C. nigoni* sequences and horizontally acquired bacterial or archaeal sequences.

Given the multiple independent origins of *Cni-hlix-1* sequences, we speculate that certain sequence components were likely derived through transposition. Supporting this, self-alignment of *Cni-hlix-1* revealed clear inverted repeats flanking exons 2-5 (from *Cni-lea-1*) and two sets of direct repeats surrounding exons 6 and 7 (potential bacterial origin), implying remnants of transposable elements (Fig. 4E). We therefore propose an evolutionary model in which local duplication of neighboring genes, i.e., *Cni-prag-2* and *Cni-otub-3*, was followed by transposition of *Cni-lea-1* sequences and horizontal acquisition of bacterial or archaeal DNAs, with subsequent local rearrangements ultimately generating the chimeric *Cni-hlix-1* gene (Fig. 4F).

To further characterize the expression pattern and function of *Cni-hlix-1*, we generated endogenously tagged versions of the gene by inserting mNeonGreen or GFP at both the C- and N-termini, as well as three internal positions between or within IDRs (Fig. S11A). We then evaluated the functionality of these tagged *Cni-hlix-1* by genotyping and assessing hybrid lethality in the IR strain ZZY10412 carrying the tagged allele on the non-introgressed Chr. IV when crossed with *C. briggsae* fathers (Fig. S11B). The absence of GFP-expressing hybrid adults, similar to the observation from crosses using ZZY10412 mothers and *C. briggsae* fathers, would be indicative of a functional *Cni-hlix-1* fusion. Notably, tagging at two sites, including at the C-terminus, completely abolished the toxin gene function, whereas the other three tagging sites retained functionality (Fig. S11B). However, neither the functional nor the nonfunctional tagged versions of *Cni-hlix-1* exhibited detectable expression in the germline or early embryos of *C. nigoni* (Fig. S11C and D). It remains unclear whether this lack of observable fluorescence results from interference with the fluorescent protein possibly due to a specific IDR of HLIX-1, or from a more restricted spatiotemporal expression pattern of the Cni-HLIX-1 protein.

### The copy number and sequence divergence of the TA gene pair among *C. nigoni* populations

Given that the TA is unique to *C. nigoni*, whether it only exist in *C. nigoni* JU1421 strain, which is the reference strain we used for hybrid crossing and phenotyping, remains unclear. To understand the evolutionary trajectory of the TA among *C. nigoni* populations, we investigated the presence/absence pattern of the gene pair among various *C. nigoni* strains we recently collected (*21*). We leveraged the recently assembled, chromosomal-level genomes of the reference strain JU1421 and eight *C. nigoni* wild isolates (*28*), and performed a phylogenetic analysis of all the *fbxn* family genes including *Cni-shls-2* (Fig. 5A). Consistent with our recent findings of highly dynamic features in both the type and number of *fbxn* genes among *C. nigoni* strains (Fig. 5A) (*28*), the *Cni-shls-2* genes exhibit considerable variations in terms of their sequences, presence/absence status and copy number (Fig. 5A). Specifically, seven out of eight *C. nigoni* wild isolates harbor *shls-2* genes, whereas strain ZF1220 lacks the gene, consistent with the fact that it possesses the fewest *fbxn* genes and lacks another member, *Cni-neib-1* (*21*). Moreover, although most Cni-SHLS-2 protein sequences are 100% identical, the sequences from the strains JU4356, EG5268, and JU2484 show noticeable divergence (Fig. 5A and Fig. S12). Besides, JU4356 and EG5268 each harbor one copy of *Cni-shls-2*, while JU2484 contains two divergent copies (Fig. 5A), whose gene models were supported by RNA-seq data (Fig. S13). Notably, three wild isolates, i.e., JU1422, YR106, and JU2617, contain multiple, identical paralogous copies of *Cni-shls-2* similar to those in JU1421, although copy numbers vary (Fig. 5A). For instance, JU1422 has three copies, YR106 has two, and JU2617 has five, as confirmed by ONT long-read sequencing (Fig. S14), indicating the highly dynamic feature of *Cni-shls-2* across populations. Interestingly, the toxin gene *Cni-hlix-1* is identified exclusively in the strains with at least two identical copies of *Cni-shls-2*, i.e., JU1421, JU1422, YR106, and JU2617 (Fig. 5A), suggesting that the presence of toxin imposes selective pressure on *Cni-shls-2* copy number.

**Figure 5.**
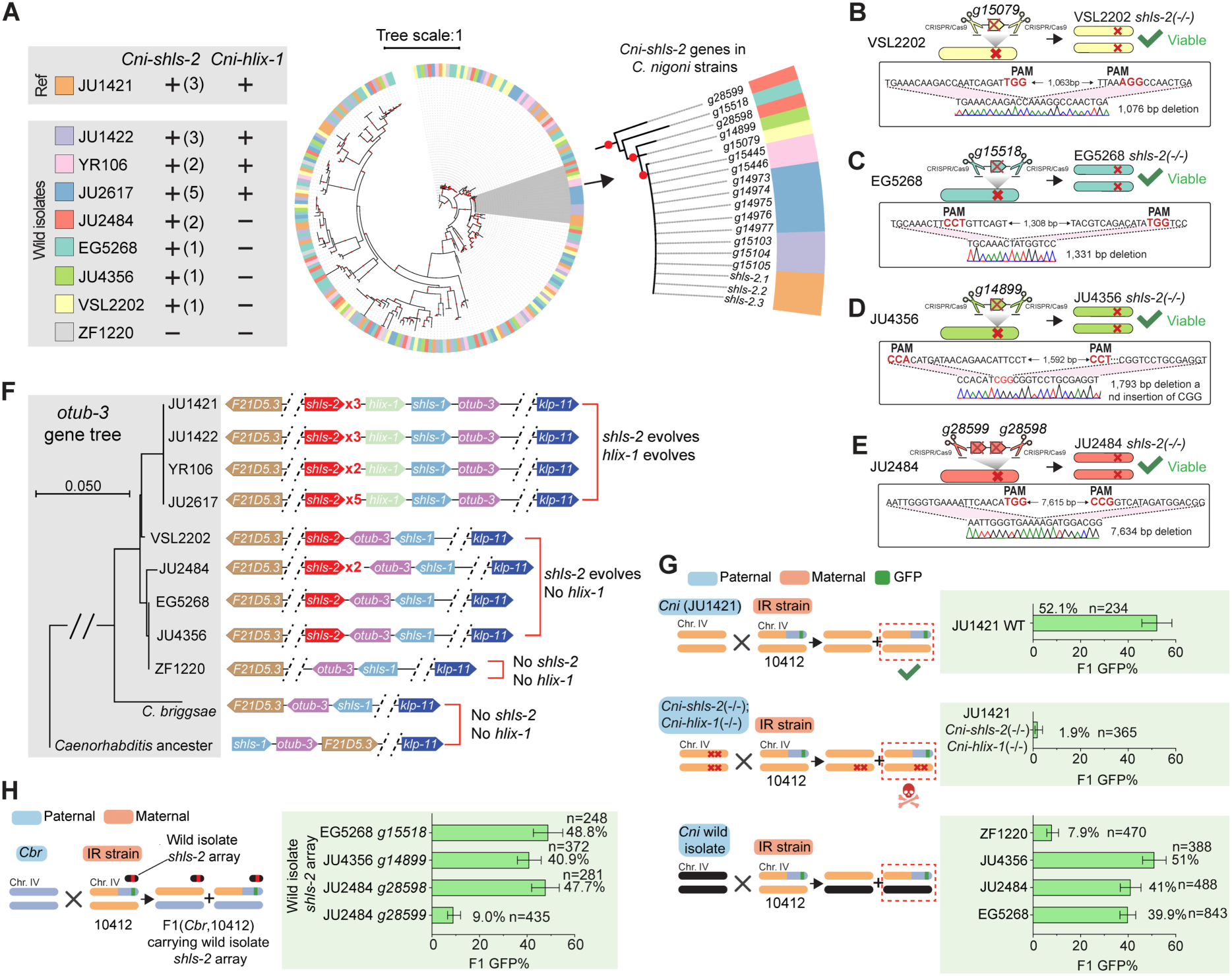
Phylogenetic analysis of *C. nigoni* populations confirms that *Cni-shls-2* evolved prior to *Cni-hlix-1*. **(A) Left:** The presence/absence status of the TA gene pair in *C. nigoni* strains. Numbers in the parentheses indicate the paralogous copy number of *Cni-shls-2*. Ref: Reference strain. **Right:** Phylogenetic tree of the *fbxn* gene family across *C. nigoni* strains. An enlarged view of the *Cni-shls-2* clade is shown aside. Red dots indicate branches with bootstrap values >80. Branch length unit: amino acid substitutions per site. **(B-E)** Schematics depicting the deletion allele of *Cni-shls-2* and the viability of the *Cni-shls-2(-/-)* mutant (top), as well as the deletion boundary (bottom) in *C. nigoni* strains VSL2202(B), EG5268 (C), JU4356 (D), and JU2484 (E). **(F) Left:** Phylogenetic tree constructed from *Cni-otub-3* protein sequences in *C. nigoni* populations and *C. briggsae* (AF16). Branch length unit: amino acid substitutions per site. **Right:** Schematic illustrating the relative positions and orientations of the TA gene pair and adjacent conserved genes including *F21D5.3*, *shls-1*, *otub-3*, and *klp-11*, in *Caenorhabditis* ancestors, *C. briggsae*, and each *C. nigoni* strain (*C. elegans* gene names are used for simplicity). **(G)** Comparison of the percentage of GFP-expressing adult F1 progeny from crosses between a WT *C. nigoni* (JU1421) father (top), a double homozygous TA gene mutant father (in JU1421; middle), or wild isolate *C. nigoni* fathers (bottom) with a ZZY10412 mother. Error bars: 95% confidence intervals. **(H)** Comparison of the percentage of GFP-expressing adult F1 progeny from crosses between a *C. briggsae* father and a ZZY10412 mother carrying an extrachromosomal array of *Cni-shls-2* derived from a *C. nigoni* wild isolate. Error bars: 95% confidence intervals.

To further validate the absence of *Cni-hlix-1* in other strains, we generated deletion mutants of *Cni-shls-2* in strains carrying only *Cni-shls-2* but lacking *Cni-hlix-1*, i.e., VSL2202, JU4356, EG5268, and JU2484 (Fig. 5B-E). As expected, these homozygous *Cni-shls-2* mutants in *C. nigoni* wild isolates were fully viable (Fig. 5B-E), indicating that *Cni-shls-2* is not essential in these strains. Given the observed presence/absence patterns of the TA gene pair across *C. nigoni* strains, i.e., both present, only *Cni-shls-2* present, or both absent, two evolutionary scenarios are possible: either the TA system evolved in the ancestral *C. nigoni*, with the toxin gene subsequently degenerating and being eliminated in some strains, or the antidote gene evolved first, with the toxin emerging subsequently in specific populations.

### The antidote gene *Cni-shls-2* predates the toxin gene *Cni-hlix-1* in *C. nigoni*

To examine which possibility is plausible, we investigated the synteny of the genomic regions carrying the TA gene pair along with their adjacent conserved genes: *F21D5.3*, *shls-1 (pgm-3)*, *otub-3*, and *klp-11* in various *C. nigoni* strains, *C. briggsae*, and other *Caenorhabditis* species. We previously showed that the consecutive arrangement in the 5’-to-3’ orientation for *Cni-shls-1*, *Cni-otub-3*, and *Cni-F21D5.3* represents an ancestral gene order in *Caenorhabditis* nematodes (Fig. 5F) (*21*). In contrast, an inversion of these three genes occurred in the common ancestor of *C. briggsae* and *C. nigoni* (*21*), as evidenced by the inverted order and orientation observed in *C. nigoni* strains that either lack the TA (ZF1220) or harbor only *Cni-shls-2* without *Cni-hlix-1* (JU4356, EG5268, JU2484, and VSL2202) as well as in *C. briggsae* (Fig. 5F and Fig. S15). Interestingly, in strains that possess both TA genes (JU1421, JU1422, YR106, and JU2617), a subsequent inversion further reversed the relative order of *Cni-shls-1* and *Cni-otub-3* (Fig. 5F and Fig. S15). These observations strongly support a model in which the antidote gene *Cni-shls-2* first evolved in the ancestral *C. nigoni*, followed by subsequent emergence of the toxin gene *Cni-hlix-1* accompanied by local rearrangement of *Cni-shls-1* and *Cni-otub-3* in only a subset of strains. Consistent with this, the coding sequences of *Cni-hlix-1* are 100% identical among all strains carrying the TA pair, whereas subtle divergence of *Cni-shls-2* sequence was observed among strains JU2484, EG5268 and JU4356 (Fig. 5F and Fig. S12), supporting a more recent evolution of the toxin relative to the antidote.

To further illustrate this possibility, we constructed a phylogenetic tree using the sequences of the adjacent, conserved gene *Cni-otub-3* and its orthologues in various *C. nigoni* strains and *Caenorhabditis* species. In agreement with our model, strains that harbor both TA components cluster together, whereas strains containing only *Cni-shls-2* or lacking the TA form a separate clade (Fig. 5F). Notably, VSL2202, the sole strain lacking the toxin but carrying *Cni-shls-2* with 100% identical protein sequence to those from the toxin-bearing strains (Fig. 5A), clusters closer to the TA-presence group (Fig. 5F), suggesting it may represent an intermediate state prior to the evolution of *Cni-hlix-1*. Collectively, these results support the notion that while the TA module is *C. nigoni*-specific, the antidote gene *Cni-shls-2* predates the toxin gene *Cni-hlix-1*, with the complete TA pair emerging only in certain *C. nigoni* lineages.

### *Cni-shls-2* can neutralize the toxin regardless of its co-existence with *Cni-hlix-1* in *C. nigoni* strains

These findings raise an intriguing question: why would the antidote gene evolve in the absence of the toxin? Although a recent study has suggested that antidote genes may initially arise as a “presuppression” mechanism to buffer against future deleterious mutations in essential genes (*13*), this scenario does not appear applicable to our system. Instead, the most parsimonious explanation for our TA module seems to be linked to host-pathogen interactions. This is because, first, F-box genes are extensively expanded in nematodes (*29*), which are widely considered to play a role in innate immune responses to various pathogenic challenges (*29–31*). Second, our results demonstrate that a portion of *Cni-hlix-1* seems to have originated from bacteria, suggesting that *Cni-shls-2* may initially evolve to target a toxic component from bacteria or the bacteria itself. Consequently, to cope with concurrent pathogens and the toxin *Cni-hlix-1*, a higher dosage of *Cni-shls-2* is needed, consistent with the observation that tandem duplications of *Cni-shls-2* were found only in those that evolved *Cni-hlix-1*. Indeed, RNA-seq data reveal that higher *Cni-shls-2* copy numbers generally correlate with elevated *Cni-shls-2* expression, although exceptions exist in certain strains (Fig. S16).

Given that *Cni-shls-2* predated *Cni-hlix-1* and *Cni-hlix-1* seems to partially originate from some bacterial components, which have taken place very recently, we reasoned that *Cni-shls-2* alleles from *C. nigoni* strains lacking *Cni-hlix-1*, which may originally target pathogens, should also be capable of detoxifying *Cni-hlix-1*. In line with this hypothesis, we found that although several amino acid differences are broadly distributed among the divergent *Cni-shls-2* alleles from JU2484, EG5268, and JU4356 compared with the identical *Cni-shls-2* alleles in other *C. nigoni* strains, almost none of these substitutions occur within the F-box-associated domain (FBA2) (Fig. S12). We subsequently assessed the functionality of these divergent *Cni-shls-2* alleles by evaluating hybrid lethality in crosses between the wild *C. nigoni* isolate fathers carrying these alleles and ZZY10412 mothers (Fig. 5G). Failure to mitigate *Cni-hlix-1* toxicity would result in a marked reduction in GFP-expressing F1 progeny, similar to the crosses between *C. briggsae* fathers with ZZY10412 mothers. As positive and negative controls, wild-type reference *C. nigoni* fathers (JU1421) and *Cni-shls-1*/*Cni-hlix-1* double homozygous mutant fathers produced approximately 52.1% and 1.9% GFP-expressing F1 progeny, respectively, when crossed with ZZY10412 mothers (Fig. 5G). As expected, all the *C. nigoni* strains, whether carrying the identical or divergent *Cni-shls-2* alleles relative to that in JU1421, rescued the hybrid lethality induced by JU1421 *Cni-hlix-1*, except the strain ZF1220, which lacks *Cni-shls-2* (Fig. 5G and Fig. S17), demonstrating that the *Cni-shls-2* alleles were able to neutralize *Cni-hlix-1* regardless of its co-existence with the toxin gene in *C. nigoni* wild isolates. To further validate this and distinguish whether the rescuing effects differ between the divergent *Cni-shls-2* alleles, we crossed *C. briggsae* fathers with ZZY10412 mothers carrying an extrachromosomal array expressing the divergent *Cni-shls-2* alleles from EG5268, JU4356 or JU2484, and then quantified the percentage of GFP-expressing adults from F1 hybrid progeny carrying the array (Fig. 5H). The extrachromosomal array of all *Cni-shls-2* alleles rescued hybrid lethality except the most divergent allele (*g28599* from JU2484), which showed marginal rescue (Fig. 5H), suggesting that this copy may be non-functional for neutralizing the toxin but might have evolved for an alternative role in the context of rapidly evolving F-box genes. Collectively, these results support sequential evolution of toxin and antidote, where initial evolution of antidote *Cni-shls-2* may be used to counteract some foreign bacterial/archaeal toxins, which also confers antagonism against subsequently internalized toxin *Cni-hlix-1*.

## Discussion

TA modules provide a compelling framework for understanding how genomic divergence leads to genetic incompatibility. While extensive research has focused on prokaryotes (*3*), recent studies indicate the potential ubiquity of these selfish systems across eukaryotes (*1*). However, few investigations have addressed a fundamental question, i.e., how and why TA modules have evolved in the first place. Like many other types of selfish genetic elements that facilitate their own propagation, TA elements typically impose detrimental effects on the fitness of hosts that fail to inherit the elements. Then why would such systems initially evolve and be maintained in the host genome? Our identification and subsequent genomic and phylogenetic analyses of the novel toxin-antidote pair, *Cni-shls-2* and *Cni-hlix-1* in the nematode *C. nigoni*, support the notion that these systems may have emerged from host-pathogen interactions. Our findings demonstrate that the antidote gene *Cni-shls-2* evolved first and the toxin gene *Cni-hlix-1* next (Fig. 5). Given that *Cni-shls-2* from strains without *Cni-hlix-1* can rescue embryonic lethality in the strains carrying both TA components (Fig. 5), we propose that initial evolution of antidote was likely driven by selective pressure imposed by pathogenic risks. The subsequent evolution of the toxin may exert selective pressure to maintain both TA components in hosts, especially when the relevant pathogens are absent, thereby ensuring immediate immunity against future exposures. Therefore, the evolution of the newly identified TA system along with many others may represent an adaptive strategy deployed during relentless host-pathogen arms races.

The identification of *Cni-shls-2* as an F-box protein-encoding gene aligns with recent findings in both plants and nematodes, where F-box proteins and, more broadly, ubiquitin ligase adaptor proteins, frequently function as antidotes in TA modules (*13, 15*). Notably, the expansion of these adaptor proteins in both nematodes and plants is widely viewed as a mechanism for innate immune defense (*29*). In nematodes, which engage continuously with diverse microbes as intermediate decomposers, such an expansion may be an adaptive necessity. Supporting this, multiple studies demonstrate that F-box genes, along with other components of ubiquitin complexes, are systematically upregulated in *C. elegans* upon exposure to various intracellular pathogens, such as the *Orsay* virus (*31*) and microsporidia (*30*). Furthermore, a differential expansion of these genes among various nematode species (*32*), particularly the pronounced expansion observed in *C. nigoni* relative to *C. briggsae* (*21, 32*), may reflect disparate pathogenic risks among different species. Supporting this, recent findings reveal a differential susceptibility to virus among different (*33*) or even within populations of *Caenorhabditis* nematodes (*34*). Given that *Cni-shls-2* belongs to the *C. nigoni*-specific F-box gene family (*fbxn* family) and is present in most *C. nigoni* strains regardless of the presence or absence of *Cni-hlix-1* (Fig. 5), it is plausible that *Cni-shls-2* initially evolved in specific *C. nigoni* populations under pathogenic selection pressures imposed by the bacteria themselves or their toxic components (Fig. 6A). In contrast, such pressures appear to be absent in *C. briggsae* and in certain populations of *C. nigoni* such as ZF1220 (Fig. 6A). Similar to recent discoveries of native microbes in wild *C. elegans* isolates (*35*), broader global sampling of *C. nigoni* may yield the types and diversity of *C. nigoni*-specific pathogens, ultimately providing direct evidence of underlying defensive genes associated with TA evolution.

**Figure 6.**
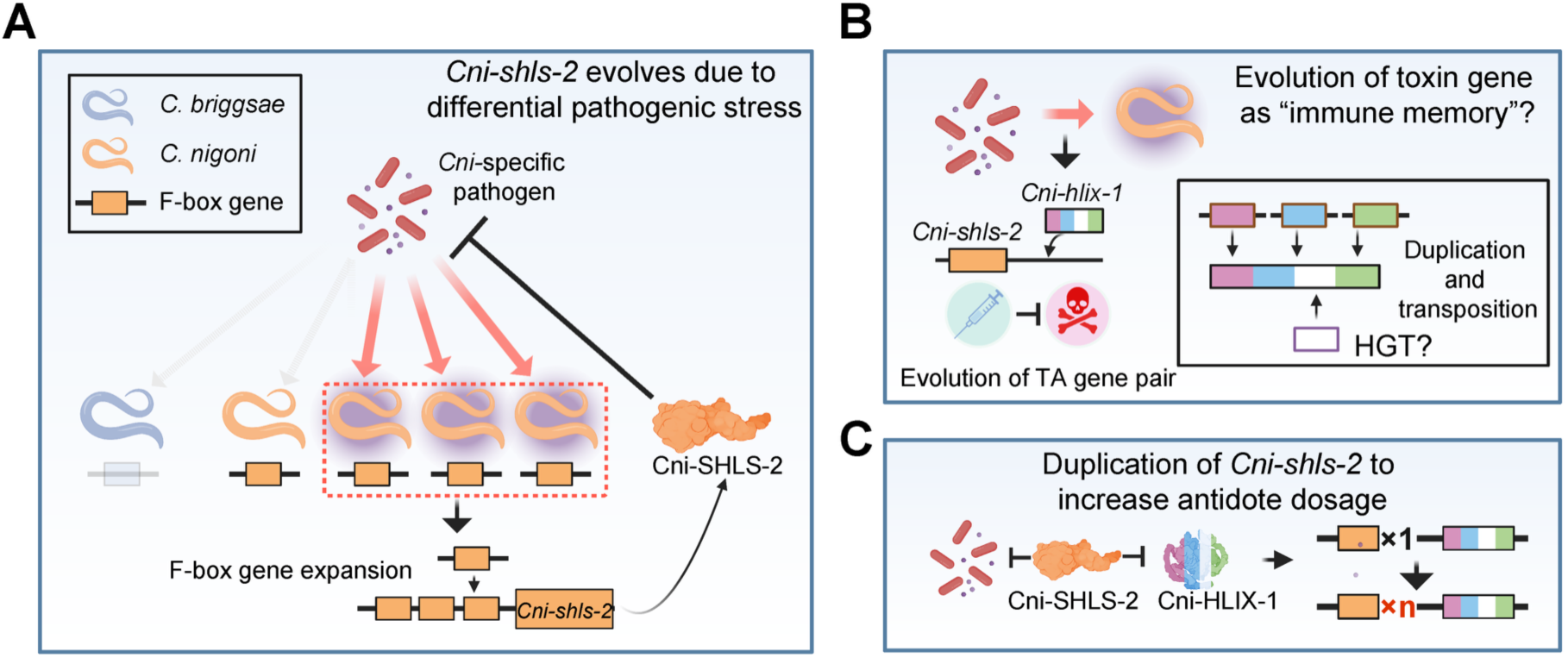
Proposed model of the evolution trajectory of the TA gene pair. **(A)** Due to habitat differences, some *C. nigoni* populations may be exposed to specific pathogens/pathogenic toxins, whereas other *C. nigoni* strains and *C. briggsae* are not. In response, *Cni-shls-2* evolves through F-box gene expansion as a defense mechanism. **(B)** To maintain or increase the immune efficacy analogous to the formation of “immune memory”, a toxin gene subsequently emerges via extensive gene reshuffling events and possibly horizontal gene transfer of pathogenic DNA sequences. **(C)** Following the evolution of *Cni-hlix-1*, *Cni-shls-2* frequently undergoes duplication, likely to increase antidote dosage for the simultaneous antagonism of both the pathogens and the toxin gene.

The evolutionary trajectory of *Cni-hlix-1* also implies an immune-related origin for the TA systems. Although direct evidence of a pathogenic origin is lacking, structural analyses reveal that a portion of the chimeric Cni-HLIX-1 toxin resembles the sequences from some bacterial/archaeal enzymes (Fig. 4). This similarity suggests that the target sequence for *Cni-shls-2* may have been domesticated via horizontal gene transfer from bacteria/archaea (Fig. 6B). Such domestication by the host from intrinsic or extrinsic parasitic agents is not uncommon (*36*). For example, domains of transposable elements are frequently co-opted for essential and novel functions in hosts (*37–39*). Subsequent gene reshuffling, incorporating fragments from conserved genes (*parg-2*, *lea-1*, and *otub-3*) (Fig. 4), likely optimized these domestication processes, refining the functional attributes of the toxin (Fig. 6B). Such steps were also observed for essential genes that evolve *de novo* through the fusion of both eukaryotic and prokaryotic sequences, as exemplified by the well-studied, essential *Drosophila* gene *oskar* (*40*). Similar to the chimeric *Cni-hlix-1*, a recent finding demonstrates extensive homology between multiple *C. tropicalis* chimeric toxin genes and the essential gene *fars-3*, implying that homologous sequences in toxin genes may arise from deleterious duplications of conserved genes (*13*). Alternatively, for *Cni-hlix-1*, the acquisition of sequences from multiple conserved genes may serve to optimize the temporal and spatial expression, as well as the functional efficacy of the toxin. For instance, since LEA-1 is a chaperone protein essential for desiccation tolerance (*41*), the intrinsically disordered regions in Cni-HLIX-1, especially those originating from *lea-1*, can either function to prevent the toxin from aggregation or increase its tolerance under stress conditions. Similarly, the deubiquitinase domain derived from *otub-3* could counteract the ubiquitination activity imposed by the antidote F-box gene *Cni-shls-2* under specific conditions, thereby fine-tuning the stability of the toxin. Supporting this, many toxin components found in *Drosophila* parasites contain deubiquitinase domains, including the OTU type (*16, 17, 42*), some of which have been shown to stabilize the toxin (*17*).

Collectively, our findings support a model in which the evolution of the toxin, following the establishment of the antidote, functions as a form of “immune memory” in response to ongoing pathogenic pressures (Fig. 6B). The formation of such TA modules, in fact, facilitates or even accelerates the fixation of the original immune-responsive gene *Cni-shls-2*, thereby enhancing the pathogen defense capabilities initially conferred by the gene. Additionally, consistent with the hypothesis, we observed tandem duplications and elevated dosage of *Cni-shls-2* exclusively in *C. nigoni* strains that harbor both toxin and antidote components (Fig. 6C). This arrangement mirrors the frequently observed tandem arrays of innate immune genes with increased gene expression, such as NLR genes in plants (*43*). A bolder speculation is that the evolution of certain TA systems may represent a transition from innate immunity to a more targeted adaptive mechanism in nematodes, analogous to CRISPR systems in bacteria (*44*).

Finally, the discovery of this novel nematode TA pair highlights the roles of immune responses in the generation of genetic incompatibilities among species and populations. The identification of two hybrid lethality suppressor genes, *Cni-shls-1* and *Cni-shls-2*, each associated with distinct incompatibility mechanisms yet both linked to immune responses involving protein degradation, illustrates that the diversification of innate immune components can fortuitously yet frequently lead to genetic conflicts and subsequent speciation in nematodes. It is intriguing that a tight linkage was observed between *Cni-shls-1* and *Cni-shls-2*, which completely blocks gene flow from *C. nigoni* to *C. briggsae* (Fig. S18). Although how such linkage originally formed remains unclear, the close physical linkage between two distinct hybrid lethality suppressor genes can lead to mutually enhanced selective pressure, ultimately resulting in a strengthened reproductive barrier (Fig. S18). With more diverse TA modules to be uncovered, we anticipate that immunity-related incompatibilities will emerge as a key driver in speciation processes across phylogenetic groups. Moreover, we speculate that at least a subset of autoimmune diseases may not necessarily be the dysfunction of the generic immune system, but rather a byproduct of specific genetic conflicts similar to those underlying *Cni-shls-1* and *Cni-shls-2* that might originally evolve for pathogen defense.

## Material and methods

### Nematode maintenance

All nematodes were cultured at 25 °C on modified Nematode Growth Medium (NGM) agar plates, with the agar concentration doubled to prevent burrowing behavior. The strains used for crossing experiments include: the reference *C. nigoni* strain JU1421; wild isolate strains of *C. nigoni*, including JU1422, EG5268, JU2484, JU2617, JU4356, VSL2202, YR106, and ZF1220 (*28*); and the reference *C. briggsae* strain AF16 with a *she-1* (V49) mutation (a spermatogenesis defect mutation used to avoid self-fertilization) (*25*).

### Generation of *Cni-shls-2* extrachromosomal array line

To zygotically express *Cni-shls-2*, the genomic sequence, along with the putative promoter and 3’ untranslated region (UTR), were PCR-amplified. The amplified fragments were subjected to gel purification prior to microinjection into the gonads of nematodes to generate a stable extrachromosomal array line. An mCherry injection marker (PZZ184) (*21*) was utilized for selection. The primers used for the amplification of *Cni-shls-2* are as follows: JU1421 *Cni-shls-2*: Forward: AGTAAGTTTGAAACAGAATTTGGGATTCGG; Reverse: AAATTATCAGTTGACCATTTCCCGTCATATGTAT. EG5268 *Cni-shls-2* (*g15518*): Forward: GGATGAGGACCATATGTCTGACG; reverse: TGAAATGGCTCAAAGTTAGGGAGCCC. JU4356 *Cni-shls-2* (*g14899*): Forward: GGCACTGACTATTCTAAGTGCAGTG; reverse: GGCTCAAAGTTAGTGAGCCCTTC. JU2484 *Cni-shls-2* (*g28598*): Forward: CTGTTTCTCTGTCAGTTTGTCCGCAC; reverse: CAGTTGACCACATGTATGCATACAGG. JU2484 *Cni-shls-2* (*g28599*): Forward: GTTTCAGTTCGATTATACCACGATCTATTCC; reverse: GTGCGGACAAACTGACAGAGAAACAG. The amplified JU1421 *Cni-shls-2* fragment corresponds to *Cni-shls-2.3*, as validated by sequencing the resulting amplicon.

### Cross setup and phenotyping

To assess the lethality ratio, at least fifty L4-stage females and seventy young adult males were placed on a 55-mm NGM plate seeded with *Escherichia coli* OP50 at the center to facilitate mating overnight. Each cross was repeated a minimum of two times. The following day, fertilized females were transferred to at least three separate 35-mm NGM plates, ensuring that each plate contained ten females. These females were allowed to lay eggs for approximately three hours before being removed. To evaluate embryonic lethality within the same species or hybrids between different species, all embryos produced during this period were counted to obtain the total egg number (n). Approximately 24 hours later, unhatched embryos were counted to determine the embryonic lethality ratio. If the percentages of hybrids with and without GFP markers were assessed, all adult progeny, both with and without GFP, were scored under a Leica M165FC fluorescence stereomicroscope two days later. The maintenance of a gene mutation as homozygous was determined by performing a 1×1 cross of the heterozygous mutant. A mutant is considered homozygotes inviable if no homozygous mutants are obtained after at least 20 plates of 1×1 crosses over three rounds of successive crosses.

### Measurement of F1 progeny lethality

F1 lethality between an introgression (IR) strain of *C. nigoni* (carrying a single *C. briggsae* introgression linked with a fluorescent marker) and *C. briggsae*, *C. nigoni*, or *C. nigoni* wild isolate strains is assessed through the absence (or near absence) of GFP-expressing adult progeny, which represents individuals lacking the *Cni-shls-2* gene or those that do not carry a functional *Cni-shls-2* gene. This measurement is valid because we observed that embryos from *C. nigoni* or its hybrids with *C. briggsae* that lack *Cni-shls-2* arrested at embryonic stages. The percentage of F1 adult worms expressing GFP is calculated by dividing the number of GFP-expressing adults by the total number of F1 adult progeny. For the crosses involving array-carrying worms, the proportion of GFP-positive worms is determined based on all adult F1, worms carrying the array, that is, expressing the mCherry fluorescent marker. Characterization of hybrid lethality between *C. briggsae* and heterozygous mutants for *Cni-shls-2* is performed by genotyping the F1 hybrid adults to determine the presence of *Cni-shls-2* wild-type/mutant alleles through PCR amplification followed by gel electrophoresis. The primers used for amplifying the wild-type *Cni-shls-2* allele are: Forward: GGTGTACTTTAGACCTGGTGACTGCG; reverse: CGCGCTCAAATAAACGAGAGGC. The primers used for amplifying Cni-shls-2 mutant alleles are the same as those for the wild-type allele: Forward: TGTAATAGAGGGGCCTCGAAAT; reverse: CAACTTTTGATCTTTAACTTGGGTATAAAAGCT.

### Characterization of arrested embryos from *Cni-shls-2 (-/-)* worms

The arrested embryos from *Cni-shls-2(-/-)* worms were obtained by collecting and observing unhatched embryos that lacked the mCherry fluorescent marker from the strain ZZY1228, which is a *Cni-shls-2(-/-)* worm carrying a *Cni-shls-2* array (ZZY1057) crossed with a homozygous mCherry::Cni-HIS-72 *C. nigoni* strain (ZZY1125). At least fifty gravid ZZY1228 females were allowed to lay eggs on a 35-mm NGM plate for approximately three hours, after which they were removed. The cell number and type of arrested embryos were characterized after at least 24 hours. We reasoned that the embryos lacking the array marker are those that do not carry the *Cni-shls-2* array, resulting in lethality due to the absence of *Cni-shls-2*. The cell number of the arrested embryos was determined by counting the number of mCherry-labeled cell nuclei. Arrested embryos were classified as “early arrest” if fewer than 28 cells were formed. Those exhibiting clear body elongation and folding were categorized as having “elongation defects,” while the remaining embryos were classified as having “gastrulation defects.”

### Validation of targeted recombination boundary

The accurate recombination sites for the IR strains ZZY10412 and ZZY10413 were verified using Oxford Nanopore Technologies (ONT) long DNA sequencing reads that span the entire recombination region, as previously described (*22*). Raw ONT reads were aligned to a manually constructed genomic region encompassing the recombination site using minimap2 (v2.17) (*45*)with default parameters.

### CRISPR/Cas9 mediated gene editing

CRISPR/Cas9-mediated genome editing was performed as previously described (*21*). Briefly, a single gRNA, consisting of both crRNA and tracrRNA, was ordered from Beijing Qingke Biotechnology Co. at a working concentration of 20 µM. The gRNA pair was mixed with Cas9 protein (glycerol-free, IDT), making the protein at a final concentration of 0.3 µg/µL, along with 10 ng/µL of injection marker solution. This mixture was kept on ice to promote ribonucleoprotein (RNP) assembly prior to microinjection. We routinely use dual gRNAs to generate large deletions spanning multiple genes, as well as smaller deletions for knockout of individual genes. For gene knock-in experiments, donor sequences were added at a final concentration of 50 ng/µL for homologous recombination. The donor sequences were amplified using a 5’ Spacer 9 (SP9) modified primer (IDT), incorporating approximately 70-nt homologous arms. The DNA amplicon was subjected to a melting process, as described by Ghanta et al., to improve knock-in efficiency (*46*). Accurate gene editing at the deletion boundaries or insertion regions was further verified by amplifying the entire sites followed by Sanger sequencing. All transgenic strains generated with corresponding gene types and gRNAs used can be found in Supplemental Table S3.

### Imaging

Images of *Cni-shls-2* homozygous mutant adult carrying a *Cni-shls-2* array (ZZY1038) were captured using a Leica M165FC fluorescence stereomicroscope. High-resolution differential interference contrast (DIC) and fluorescence imaging were performed on a Leica Stellaris confocal system, using a 60× water-immersion objective. To visualize GFP::Cni-HLIX-1 and Cni-HLIX-1::mNeonGreen adult females, approximately one day old young adults were collected and immobilized with 10 mM sodium azide. Whole-worm images were obtained utilizing the tile-scan mode. For embryo imaging, 20-30 gravid females were dissected, and the embryos were mounted on slides containing a pre-dried 0.5 mg/mL poly-L-lysine pad to adhere them to the slides. Fluorescence signals were recorded using 488-nm (GFP) and 561-nm (mCherry) lasers, set to 3% power with a detector gain of 300. All images (frame size: 712 × 512 pixels) were acquired at a scan rate of 100 Hz and with a 2.5-pinhole setting.

### Multiple sequence alignment and phylogenetic tree building

The *fbxn* gene protein sequence alignments were performed using MAFFT (v7.525) (*47*) with default settings, while the phylogenetic trees were constructed using iqTree (v2.1.2) (*48*) with the parameters “-m TEST -bb 1000 -alrt 1000.” The resulting trees were visualized and adapted from iTOL (*49*). Multiple protein sequence alignments of *C. nigoni fbxn* genes and *Cni-shls-2* paralogs in *C. nigoni* wild isolates, as well as genomic DNA sequence alignments of *shls-2.1*, *shls-2.2*, and *shls-2.3*, were conducted using MEGA XI (*50*) with the MUSCLE algorithm, and the results were visualized using Jalview (v2.11.3.2) (*51*). Self-alignments of genomic regions containing *Cni-shls-2*, *Cni-hlix-1* genomic sequences, and alignments of genomic regions of *Cni-hlix-1* with the three homologous genes were performed using BLASTn (2.7.1) (*52*). The *Otub-3* gene tree was constructed using protein sequences from all nine *C. nigoni* strains, *C. briggsae* (AF16), and *C. elegans* (N2).

### Remote homology and protein structure search

Remote homology searches of *Cni-hlix-1* sequences were performed using HHblits (*53*). Its protein domains were predicted using InterProScan (*54*). Predicted Cni-HLIX-1 structures were generated using the AlphaFold Server (*55*). The resulting models were examined in PyMOL2 (v3.0.3). Protein structural alignment was performed using the DALI protein structure comparison server (*56*).

### Validation of the genomic regions containing the TA pair

To confirm the presence or absence of the TA gene pair, ONT long reads were mapped to the chromosome-level genomes of each *C. nigoni* strain using minimap2 (v2.17). To validate the paralog number of *Cni-shls-2*, only reads spanning the entire region containing all *Cni-shls-2* paralogs were retained. The order and orientation of the TA gene pair and its adjacent conserved genes were verified by inspecting the genome assemblies supported by ONT long-read alignments. Notably, the original assemblies of the genomic regions surrounding the TA gene pair in strains EG5268, JU4356, and JU2484 appeared inaccurate, likely because their genomes were assembled using JU1421 as a reference. The correct genome sequences of local regions and corresponding gene annotations were manually curated, which is further validated by ONT long reads.

### RNA-seq data analysis

ONT native RNA-seq long reads and Illumina RNA-seq reads were mapped using minimap2 (v2.17) and STAR (v2.7.0b) (*57*), respectively. Mapped reads were visualized in IGV (*58*). To measure the expression levels of *Cni-shls-2* genes in *C. nigoni* reference and wild isolate strains, featureCounts (v2.20.0) (*59*) was used to obtain raw read counts, and DESeq2 (v1.40.2) was used to calculate normalized expression values. To ensure valid comparisons, only one-to-one orthologs and the *Cni-shls-2* genes were retained for normalization. Expression values of the multiple *Cni-shls-2* paralogs within a *C. nigoni* strain, if present, were combined and treated as the expression of *Cni-shls-2* gene.

### Statistical analysis and schematic diagram drawing

Error bars for all GFP-expressing adult worms, embryonic lethality and hatching rates represent the 95% confidence intervals of the ratios, calculated using the Agresti-Coull method. The error bar for the cell number of arrested *Cni-shls-2* homozygous mutants represents the standard error of the mean (SEM). The significance of the difference (*P* value) was calculated with the Fisher’s exact test. All schematic diagrams were created in BioRender and further modified using Adobe Illustrator.

### Genome assembly and sequencing data used

The *C. nigoni* (CN3) and *C. briggsae* (CB5) reference genome were retrieved from NCBI BioProject PRJNA917437, and their annotation files were obtained from Github (https://github.com/PikaPatch/CB-CN) (*22*). Genomes of *C. nigoni* wild isolates were obtained from the European Nucleotide Archive under accession number PRJEB100562, and annotation files were retrieved from Github (github.com/PikaPatch/F-box/tree/main/genome) (*28*). ONT DNA-seq reads and NGS RNA-seq reads for *C. nigoni* strains were obtained from NCBI BioProject PRJNA1219697 (*28*). ONT native RNA-seq reads from *C. nigoni* embryos were retrieved from NCBI BioProject PRJNA1068783 (*21*). Adult female RNA-seq reads were obtained from NCBI BioProject PRJNA659254 (*60*). ONT reads spanning the targeted recombination boundaries of ZZY10412 and ZZY10413 are obtained from NCBI BioProject PRJNA947947 (*21*).

## Acknowledgements

We thank Marie-Anne Félix, Luc Barre, Lise Frézal, Mirko Francesconi, Rémy Froissart, Takao Inoue, Varsha Singh, and Henrique Teotónio for providing *C. nigoni* wild isolate strains. We also thank Olivia Xuan Wan for the insightful discussions. We thank Mr. Chung Wai Shing for logistical support and members of Z.Z.’s laboratory for helpful discussions and comments. This work was supported by General Research Funds from the Hong Kong Research Grants Council (HKBU12100825, HKBU12101522, HKBU12101323, HKBU12100024), the Hong Kong Innovation and Technology Fund (GHP/176/21SZ), the Environment and Conservation Fund (2023-160) from the Hong Kong Environmental Protection Department, and Seed Funding for Collaborative Research Grants (RC-SFCRG/24-25/R1/SCI/01) from Hong Kong Baptist University to Z.Z. The work was also supported by the Young Scientists Fund from National Natural Science Foundation of China (32400491), and the General Research Fund from the Hong Kong Research Grants Council (HKBU12100925) to X.D. This study was supported by the Wu Jieh Yee Institute of Translational Chinese Medicine Research, Hong Kong Baptist University.

## Author contributions

X.D. and Z.Z. conceived the project. X.D., M.Y., Z.J. and Y.P. performed experiments. X.D. analyzed data. Z.Z. coordinated the project, provided guidance and resources. X.D. and Z.Z. wrote the manuscript.

