## Supplementary Figures 1-18 for "Sequential evolution of antidote and toxin links genetic incompatibility with immune responses"

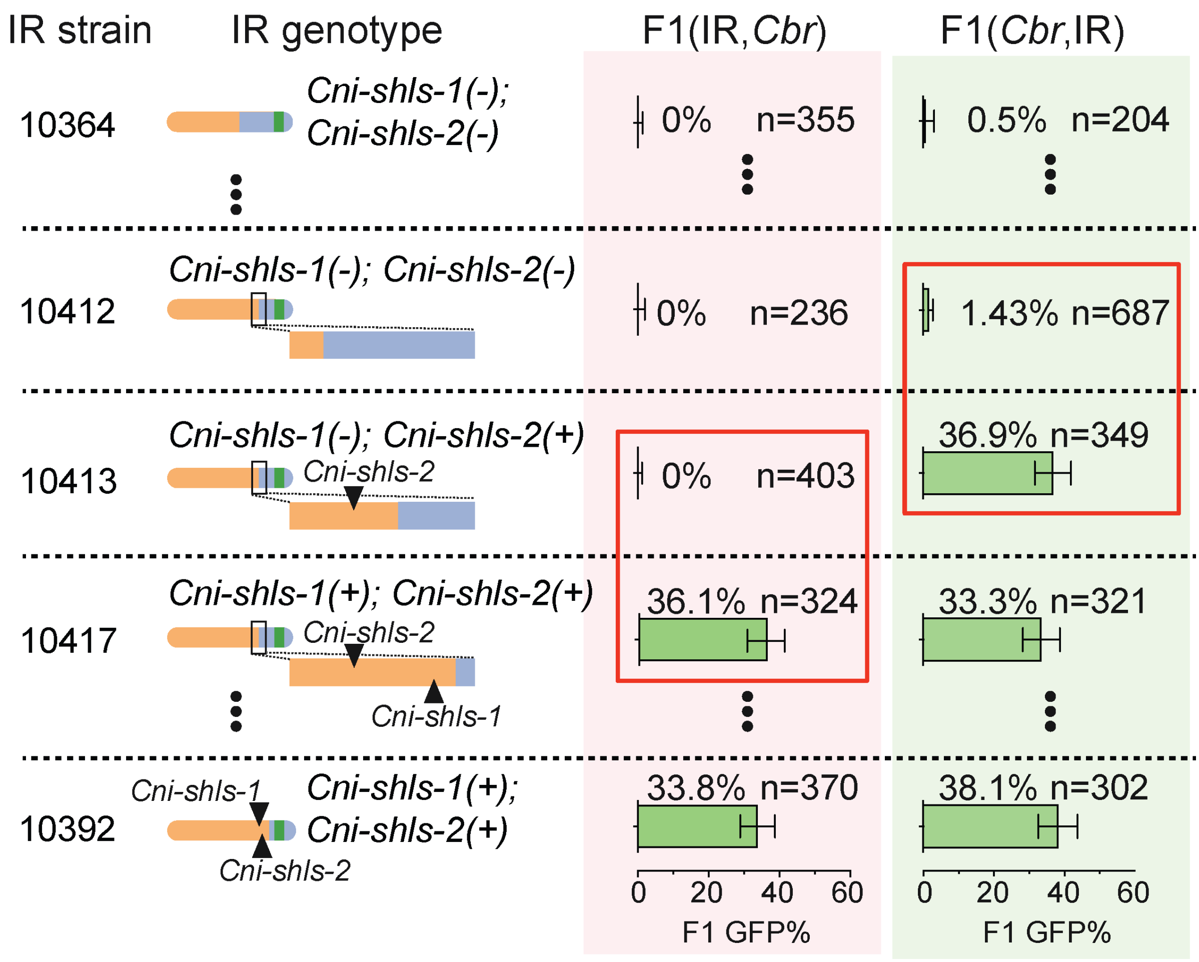


**Figure S1. Fine-mapping of the *Cni-shls-2* genetic locus using IR strains with variable *C. briggsae* introgression sizes.**

Percentages of GFP-expressing hybrid F1 adults from crosses between *C. briggsae* mothers and IR fathers (shaded with light pink) and the reciprocal crosses (shaded with light green) are shown. IR strain names and their corresponding genotypes (i.e., Cni-Chr. IV with various *C. briggsae* introgression sizes) are indicated. Note that the distinct intervals containing *Cni-shls-1* and *Cni-shls-2*, determined by two different IR strain pairs, are highlighted by red rectangles.

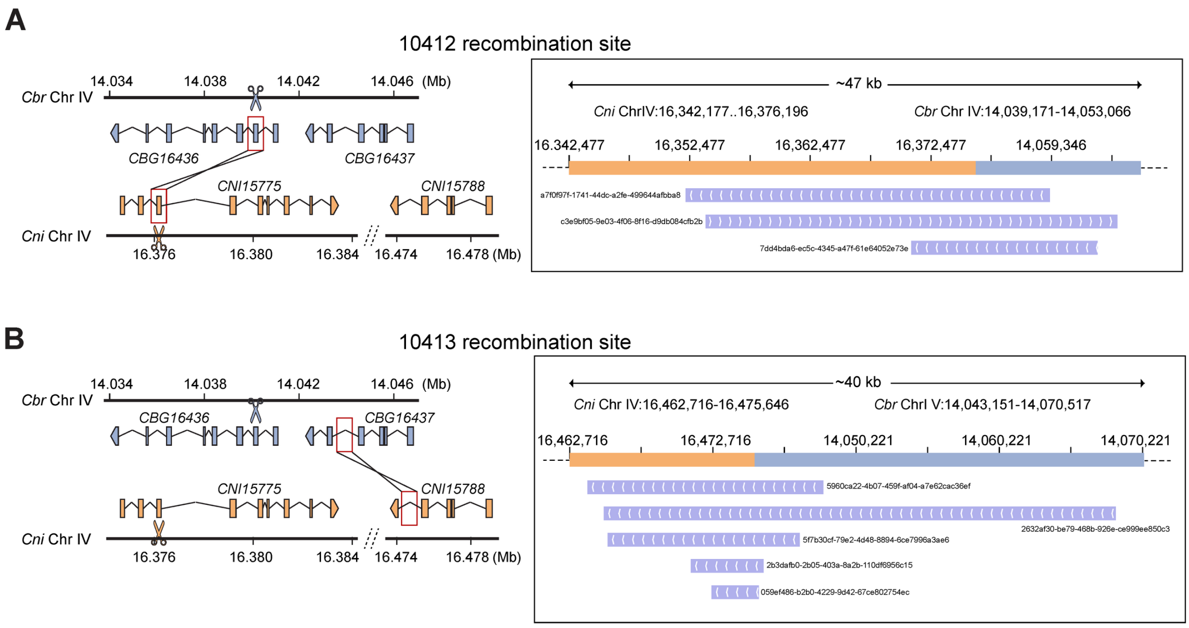


**Figure S2. Validation of recombination boundaries in key IR strains (ZZY10412 and ZZY10413) using boundary-spanning ONT (Oxford Nanopore Technology) long reads.**
**(A-B)** Genomic positions and predicted gene models for recombination sites in C. briggsae and C. nigoni (left), and the mapping tracks of ONT long reads across the recombination boundaries (right) for ZZY10412 (A) and ZZY10413 (B). Recombined sites are highlighted with red rectangles, and CRISPR/Cas9 cutting sites are marked with scissor symbols. The names of the recombination-spanning reads are indicated.

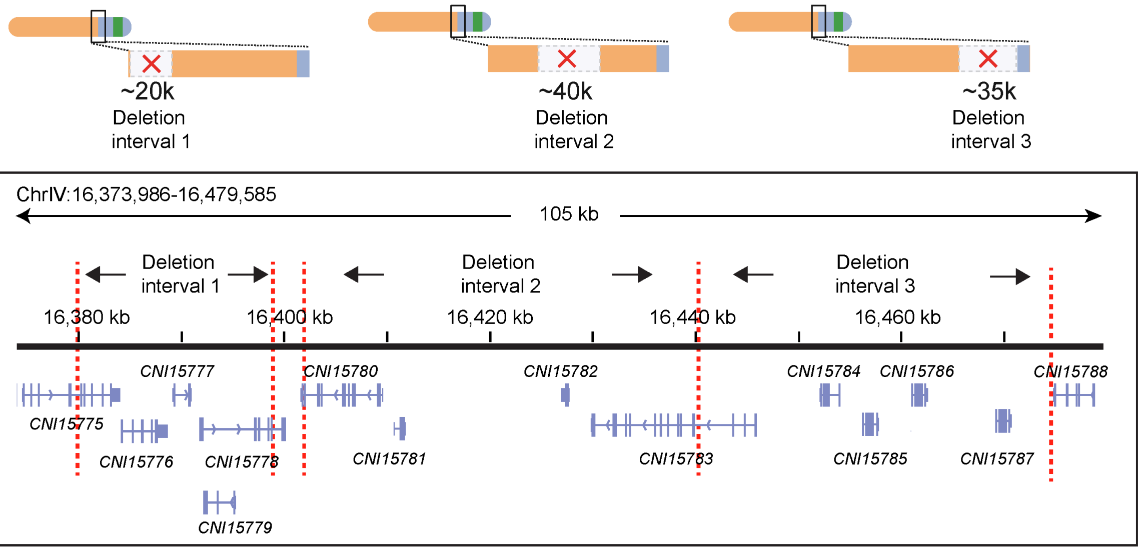


**Figure S3. Genomic positions and predicted gene models within the ~100 kb candidate interval for *Cni-shls-2*.**
Red dashed lines delimited the three large deletion intervals within the ~100 kb region, as shown in Fig. 1G.

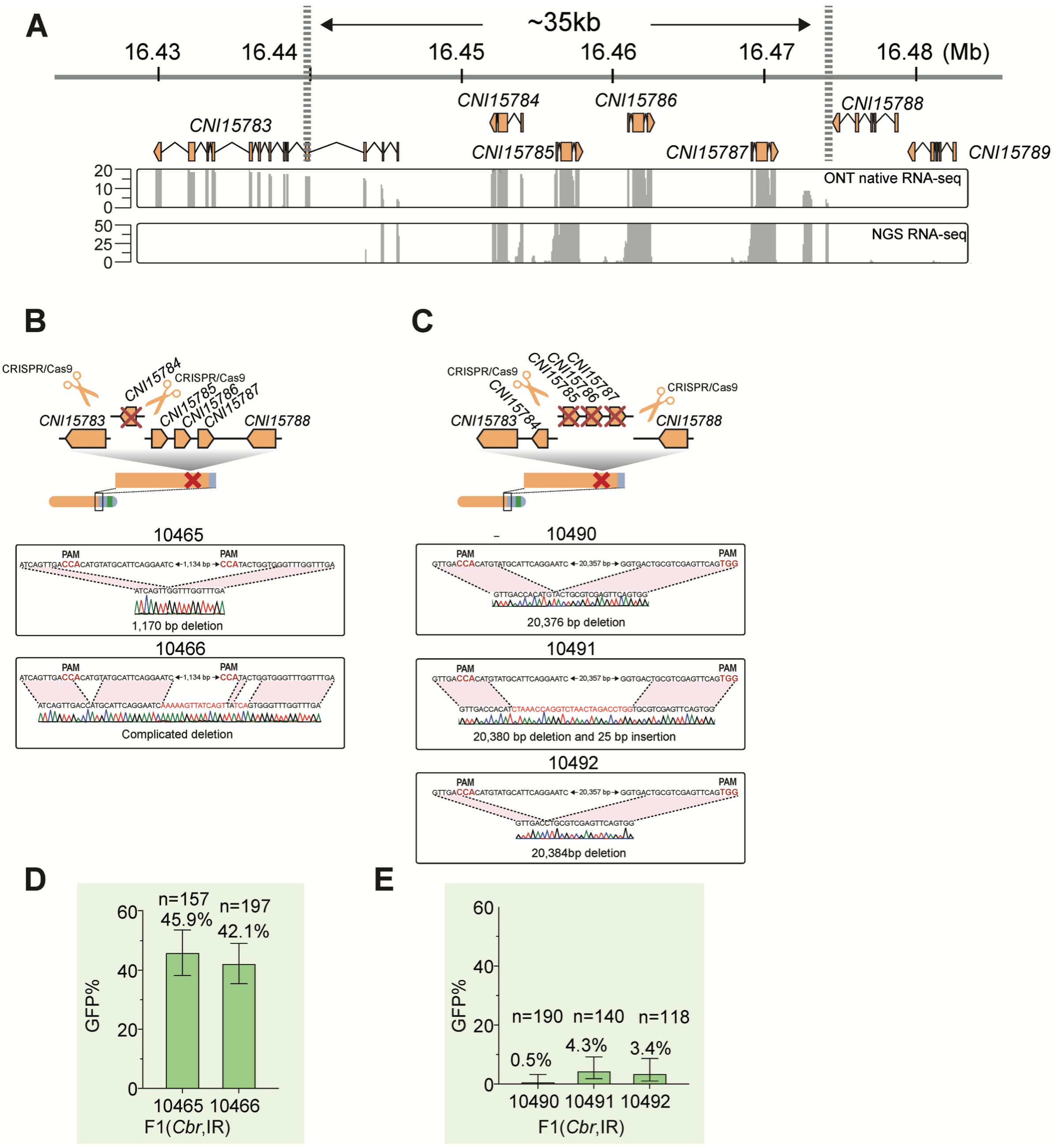


**Figure S4. *Cni-shls-2* maps to three identical genes within the ~35-kb candidate interval.** **(A)** Genomic positions and predicted gene models within the ~35-kb candidate interval for *Cni-shls-2*. RNA-seq read coverages (ONT and NGS) from *C. nigoni* embryos are displayed below.

**(B-C)** Schematics illustrating the generation of gene null mutations via CRISPR/Cas9-mediated deletion (top) and the corresponding sequence details of the deletion boundaries for *CNI15784* (B) or the three genes i.e., *CNI15785*, *CNI15786*, and *CNI15787* simultaneously (C).

**(D-E)** Bar plots showing the percentage of GFP-expressing adult F1 progeny from crosses between a *C. briggsae* father and IR mothers carrying deletions of *CNI15784* (D), or of the three genes i.e., *CNI15785*, *CNI15786*, and *CNI15787* simultaneously (E).

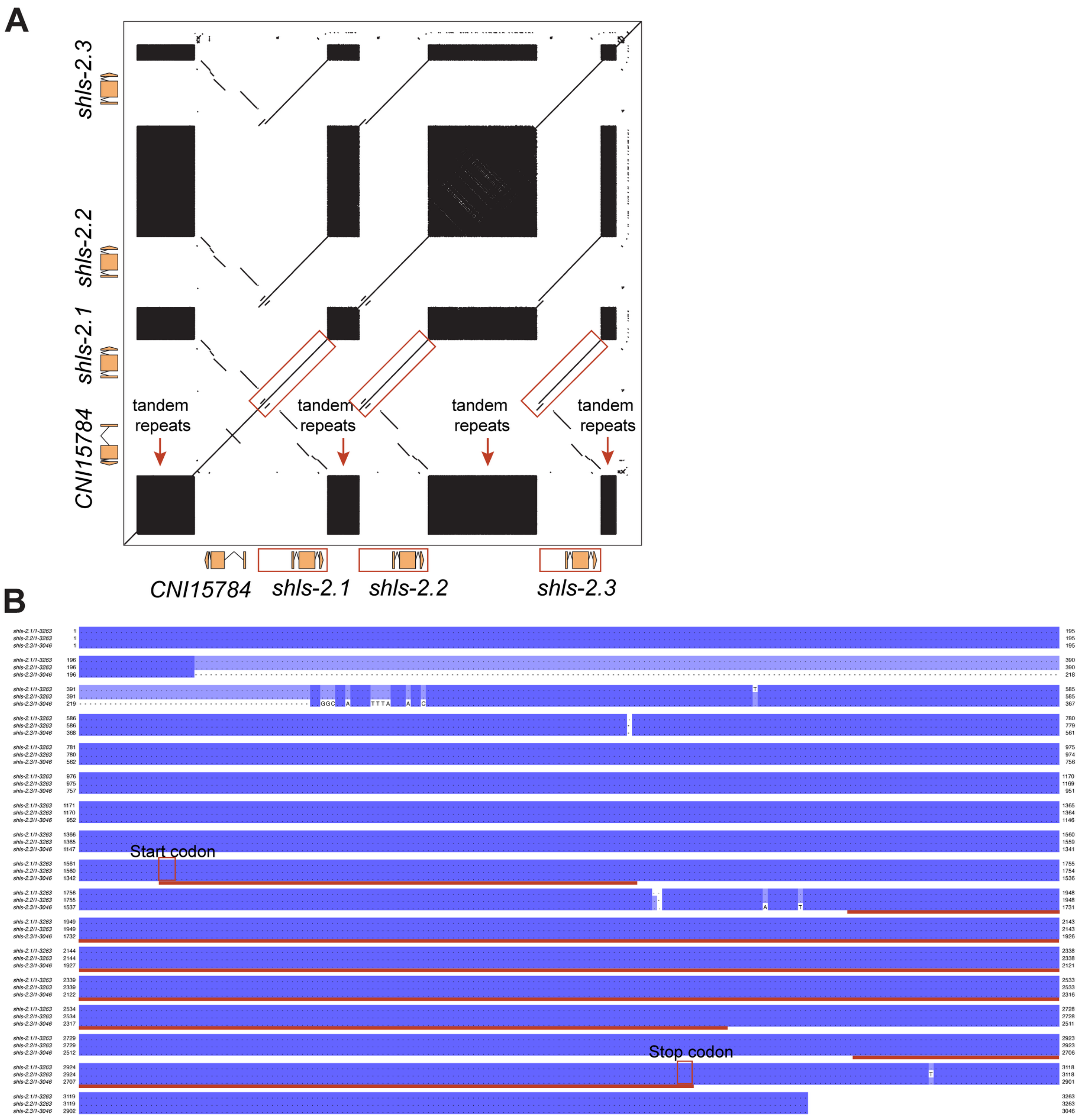


**Figure S5. The *Cni-shls-2* genetic locus comprises three identical genes.**

**(A)** Dot plot displaying the self-alignment of genomic regions harboring *fbxn-2*, *Cni-shls-2.1 (fbxn-3), Cni-shls-2.2 (fbxn-4)*, and *Cni-shls-2.3 (fbxn-5)*. Nearly identical genomic segments, including the upstream and downstream regions of the *Cni-shls-2* paralogs are highlighted with red rectangles. Note that numerous tandem repeats of the same type are present in the intergenic regions.

**(B)** Multiple sequence alignment of the genomic sequences of *Cni-shls-2.1*, *Cni-shls-2.2*, and *Cni-shls-2.3*. Identical sequences are shaded in dark purple, and the coding sequence regions are delineated by red lines. Note that, aside from minor indels outside the coding regions, an ~210-bp deletion is exclusively present in *Cni-shls-2.3*.

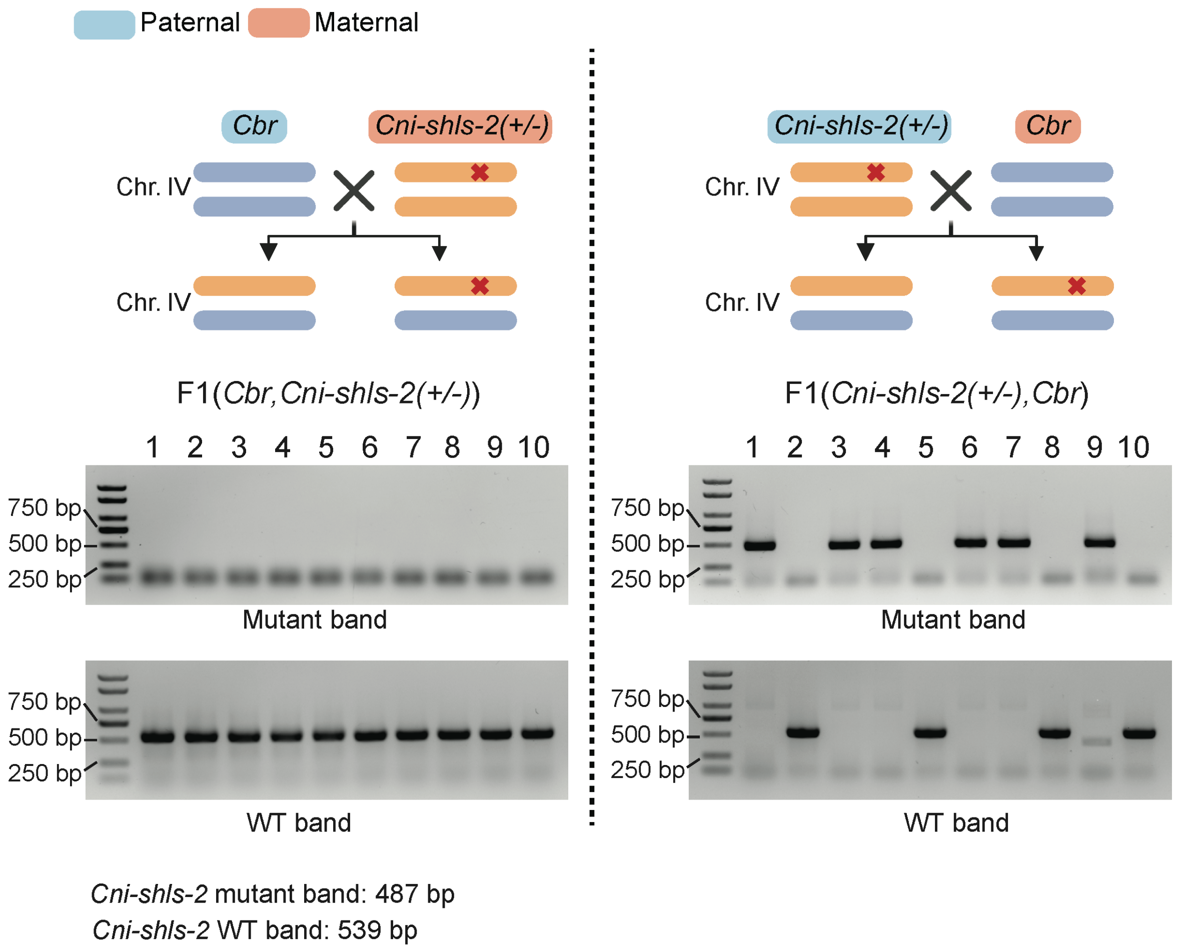


**Figure S6. Absence of *Cni-shls-2* results in lethality of hybrid progeny when *C. nigoni* serves as the mother, but not as the father.**

**Top:** Schematic of the crossing setup between *C. briggsae* and *Cni-shls-2(+/-)* mutants using reciprocal crosses.

**Bottom:** Representative gel images showing the presence and absence of the *Cni-shls-2* mutant (deletion) band and the wild-type band in 10 randomly selected F1 hybrids from crosses between a *C. briggsae* father and a *Cni-shls-2(+/-)* mother (left) or between a *C. briggsae* mother and a *Cni-shls-2(+/-)* father (right).

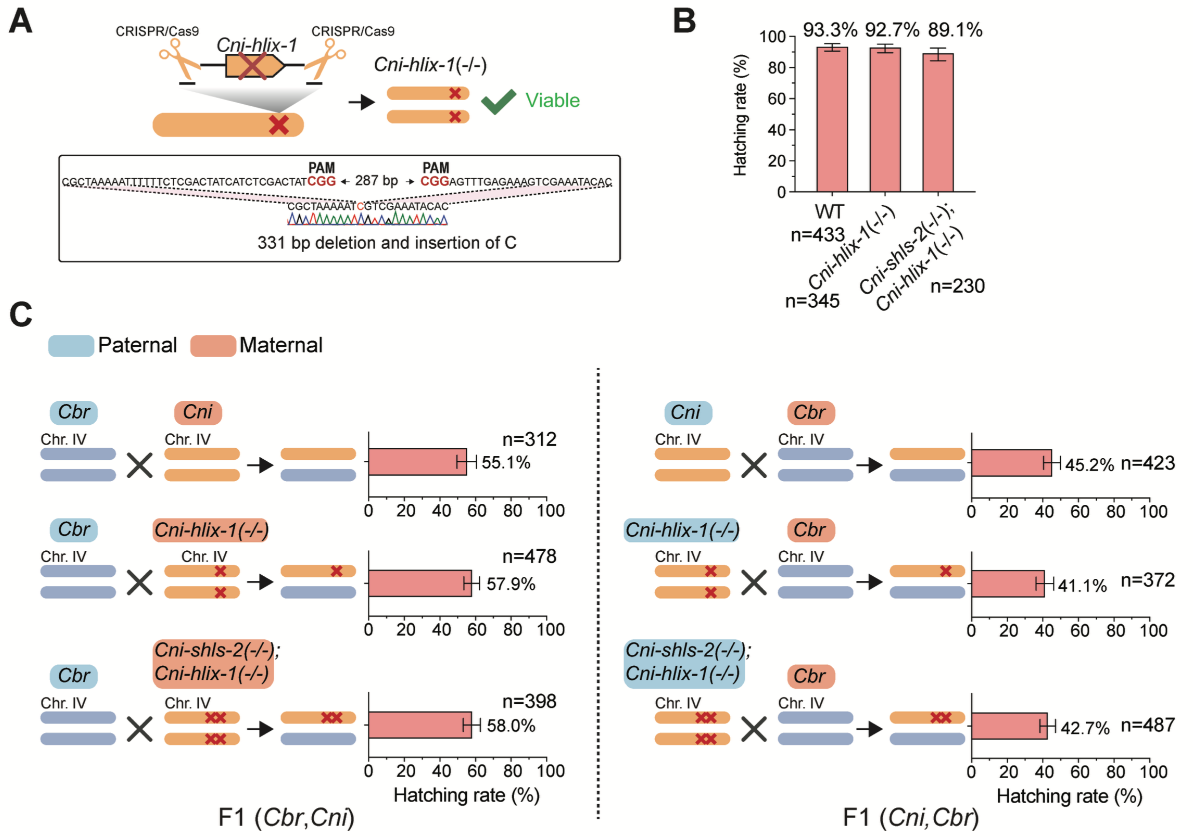


**Figure S7. Neither the absence of *Cni-hlix-1* nor simultaneous deletion of the TA gene pair significantly affects the viability of *C. nigoni* or its hybrids with *C. briggsae*.**

**(A)** Schematic representation and deletion junction sequence details of a *Cni-hlix-1* mutant allele generated by CRISPR/Cas9. Note that the *Cni-hlix-1(-/-)* mutant is viable.

**(B)** Bar plot comparing the hatching percentage of embryos among wild-type *C. nigoni*, *Cni-hlix-1(-/-)* mutants, and *Cni-shls-2(-/-)*; *Cni-hlix-1(-/-)* double mutants. No significant differences were observed among these groups.

**(C)** Schematic of the crossing setup and corresponding bar plots comparing the hatching percentage of embryos for hybrids derived from crosses with *C. briggsae* fathers and either a wild-type *C. nigoni* mother (top), a *Cni-hlix-1(-/-)* mutant mother (middle), or a *Cni-shls-2(-/-)* *Cni-hlix-1(-/-)* double mutant mother (bottom) (left), as well as the reciprocal crosses (right). No significant differences were observed among the crosses for each crossing direction.

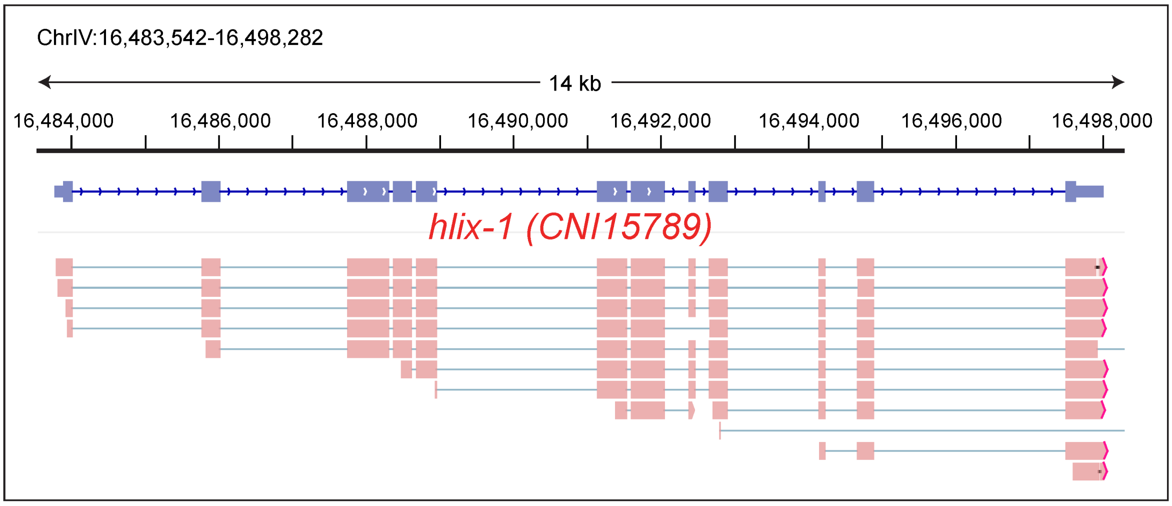


**Figure S8. ONT native RNA-seq read mapping supports the predicted gene model of *Cni-hlix-1*.**

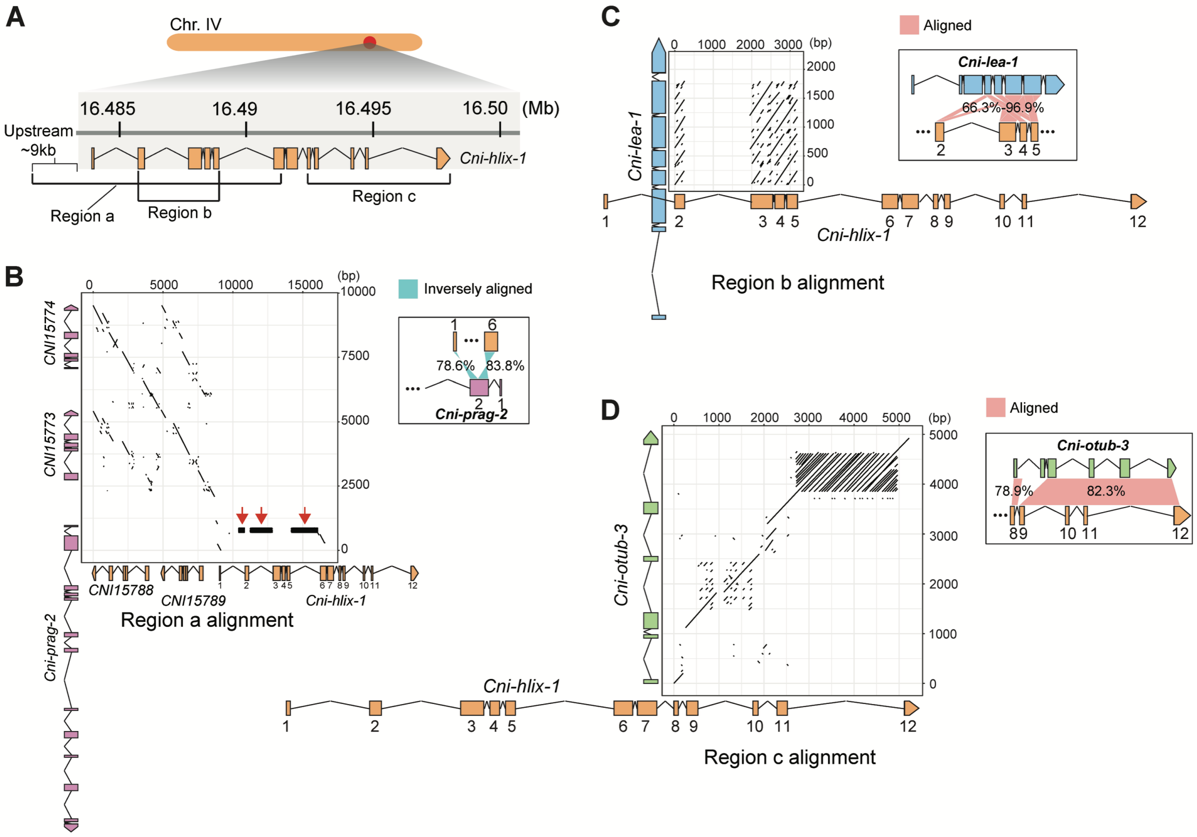


**Figure S9. *Cni-hlix-1* is a chimeric gene resulting from extensive gene reshuffling events.** **(A)** Schematic representation and genomic position of three regions of *Cni-hlix-1*. Region a: ~9 kb upstream region (including genes *CNI15788* and *CNI15789*) and exons 1-6 (region a). Region b: exons 2-5. Region c: exons 8-12.

**(B-D)** Dot plots showing the alignments between specific regions of *Cni-hlix-1* as shown in A and their corresponding genes that share homology. Region a of *Cni-hlix-1* is aligned with the first two exons of *Cni-prag-2* and its two upstream genes, *CNI15773* and *CNI15774* (B). Region b of *Cni-hlix-1* is aligned with *Cni-lea-1* (C). Region c of *Cni-hlix-1* is aligned with *Cni-otub-3* (D). Note that the introns 1, 2 and 5 of *Cni-hlix-1*, which consist of abundant tandem repeats and share homology with introns 1 of *Cni-prag-2*, are highlighted with red arrowheads. Gene models, shown to scale, are presented alongside the dot plots. Schematics of homology between different parts of *Cni-hlix-1* with the three genes (as shown in Fig. 4A) are also shown aside.

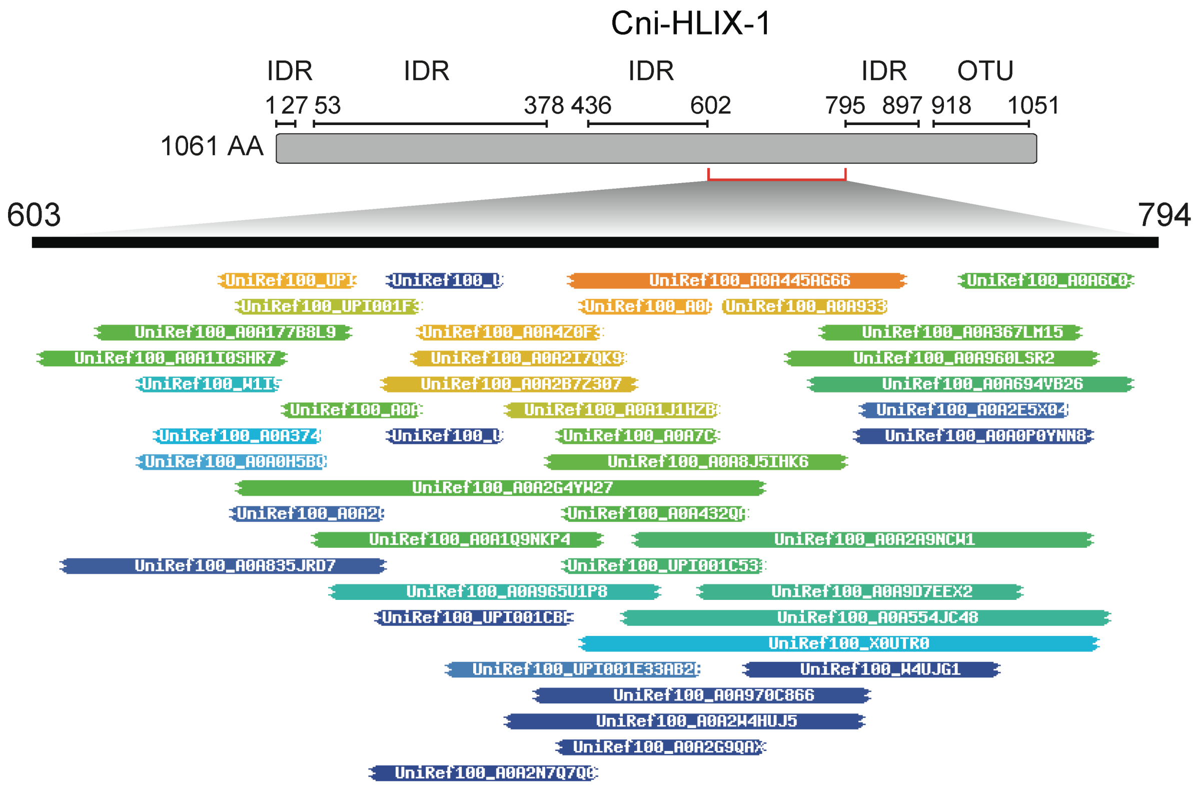


**Figure S10. Graphical representation of the hit lengths and positions for all HHblits results (excluding those against *C. nigoni*) for the approximately 191-amino acid segment in Cni-HLIX-1 that lacks homology to other nematode genes.**
Note that most hits are relatively short.

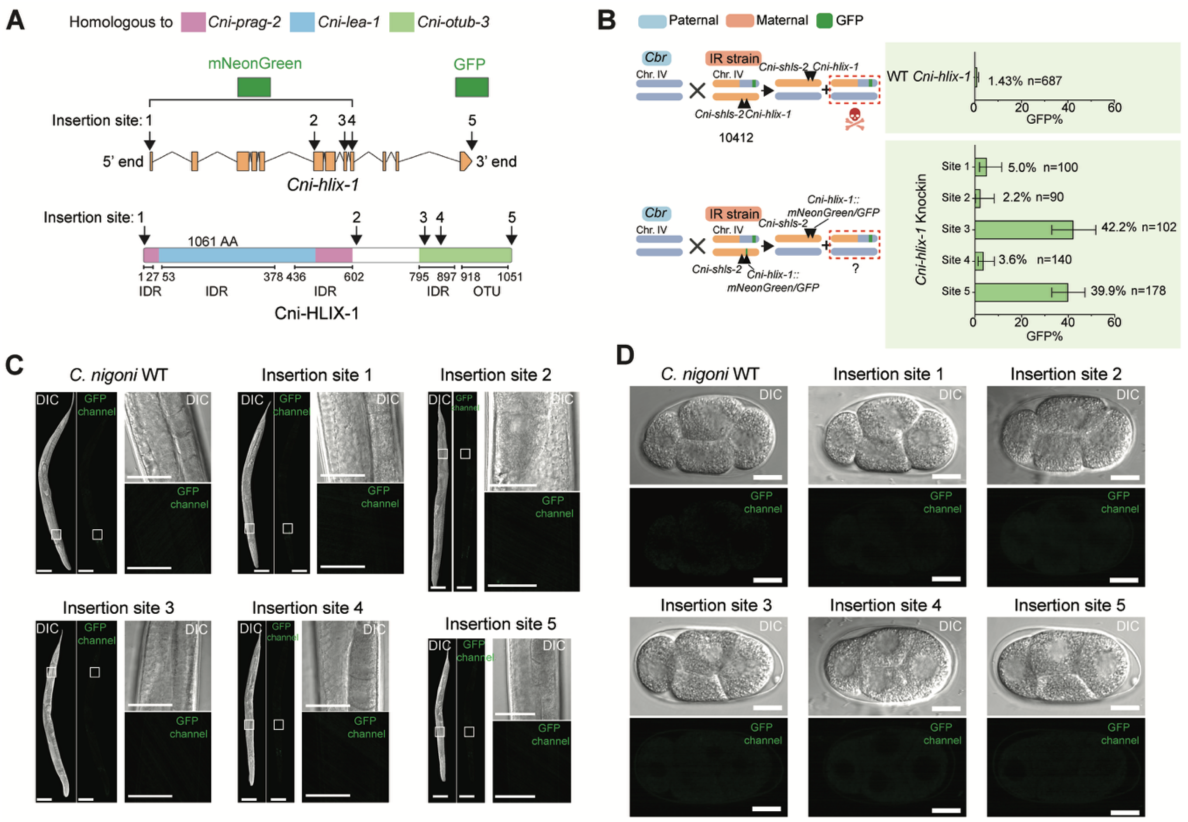

**Figure S11. No clear signals were observed for *Cni-hlix-1* endogenously tagged with fluorescent markers at various sites.**

**(A)** Schematics showing the endogenous insertion sites for mNeonGreen or GFP within the genomic sequence of *Cni-hlix-1* and the corresponding positions in the Cni-HLIX-1 protein. The segment of the *Cni-hlix-1* coding sequence homologous to *Cni-lea-1*, *Cni-prag-2*, and *Cni-otub-3*, along with the predicted protein domains of Cni-HLIX-1, is differentially color-coded as in Fig. 3G.

**(B)** Bar plots comparing the percentage of GFP-expressing hybrid adults from crosses using *C. briggsae* fathers with either a heterozygous IR strain carrying the wild-type *Cni-hlix-1* (ZZY10412, top) or a strain carrying *Cni-hlix-1*::mNeonGreen or *Cni-hlix-1::*GFP at the five positions indicated in A (bottom). Error bars: 95% confidence intervals.

**(C-D)** Comparison of DIC and fluorescent images for an adult female (C) and a 4-cell stage embryo (D) from wild-type *C. nigoni* or transgenic *C. nigoni* carrying the endogenous mNeonGreen or GFP tag at the five sites shown in A. Insets provide enlarged views of germline regions for each adult female. Scale bars: 200 µm (whole worms), 30 µm (insets), and 10 µm (embryos).

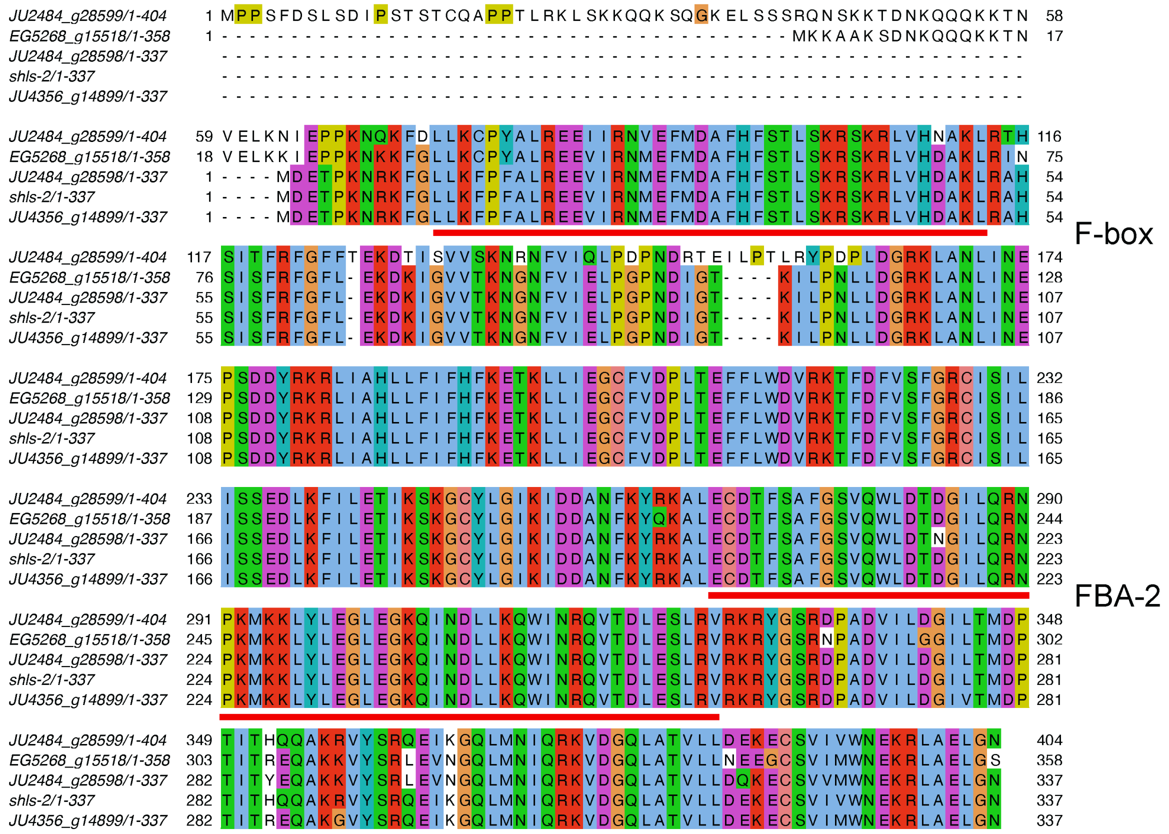

**Figure S12. Multiple sequence alignment of JU1421 Cni-SHLS-2 and divergent Cni-SHLS-2 proteins in *C. nigoni* strains**

All the *C. nigoni* strains with a Cni-SHLS-2 sequence identical to that of JU1421 are excluded. Note that the F-box and FBA-2 domains (highlighted in red) are nearly 100% identical across the strains.

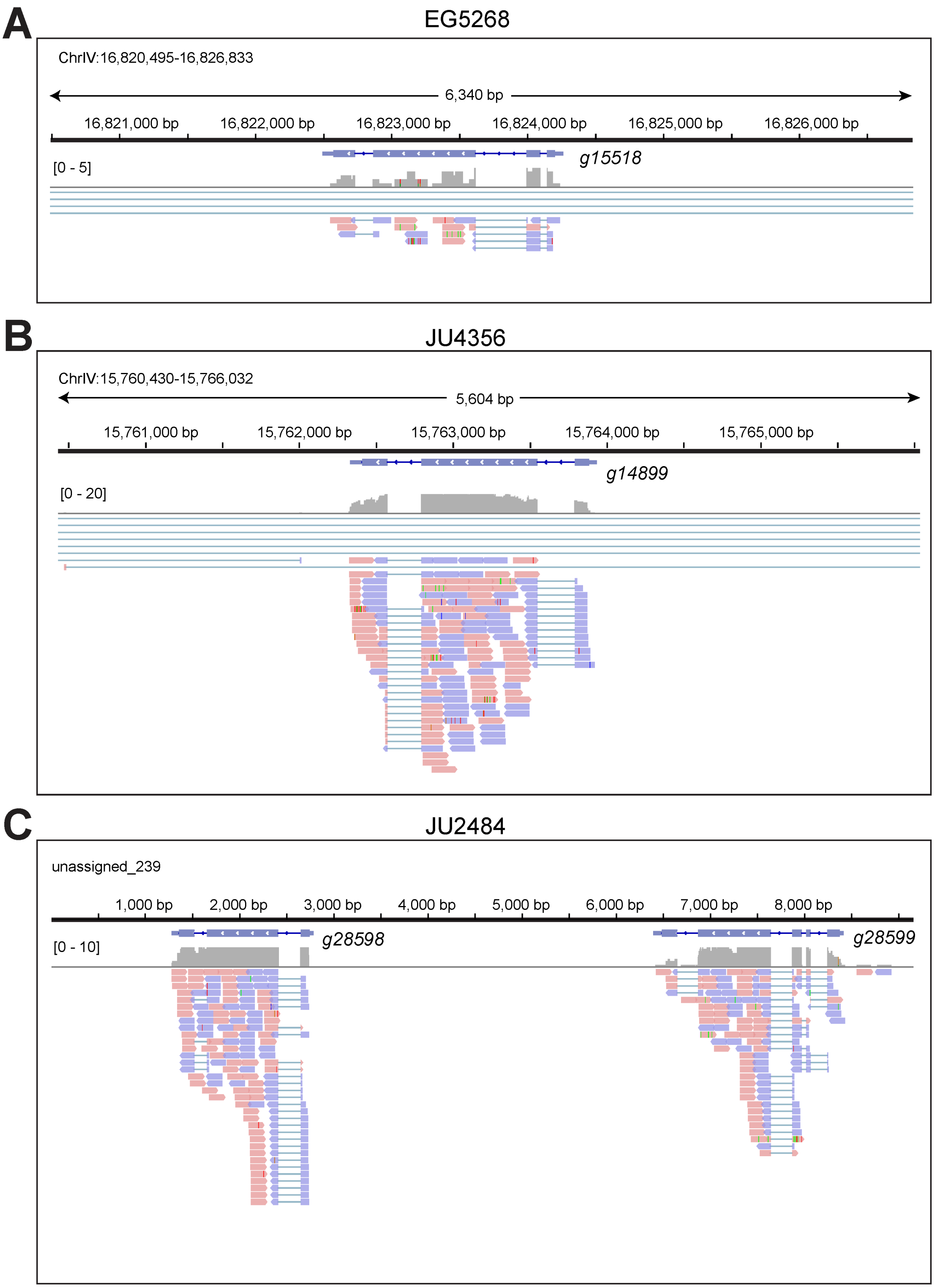


**Figure S13. RNA-seq reads confirm the gene model of the diverged *Cni-shls-2* in *C. nigoni* wild isolate strains.**

**(A-C)** Genomic positions, predicted gene models, and aligned RNA-seq reads for *Cni-shls-2* are shown for isolates EG5268 (A), JU4356 (B), and JU2484 (C). The mapped reads are colored according to their mapped strand.

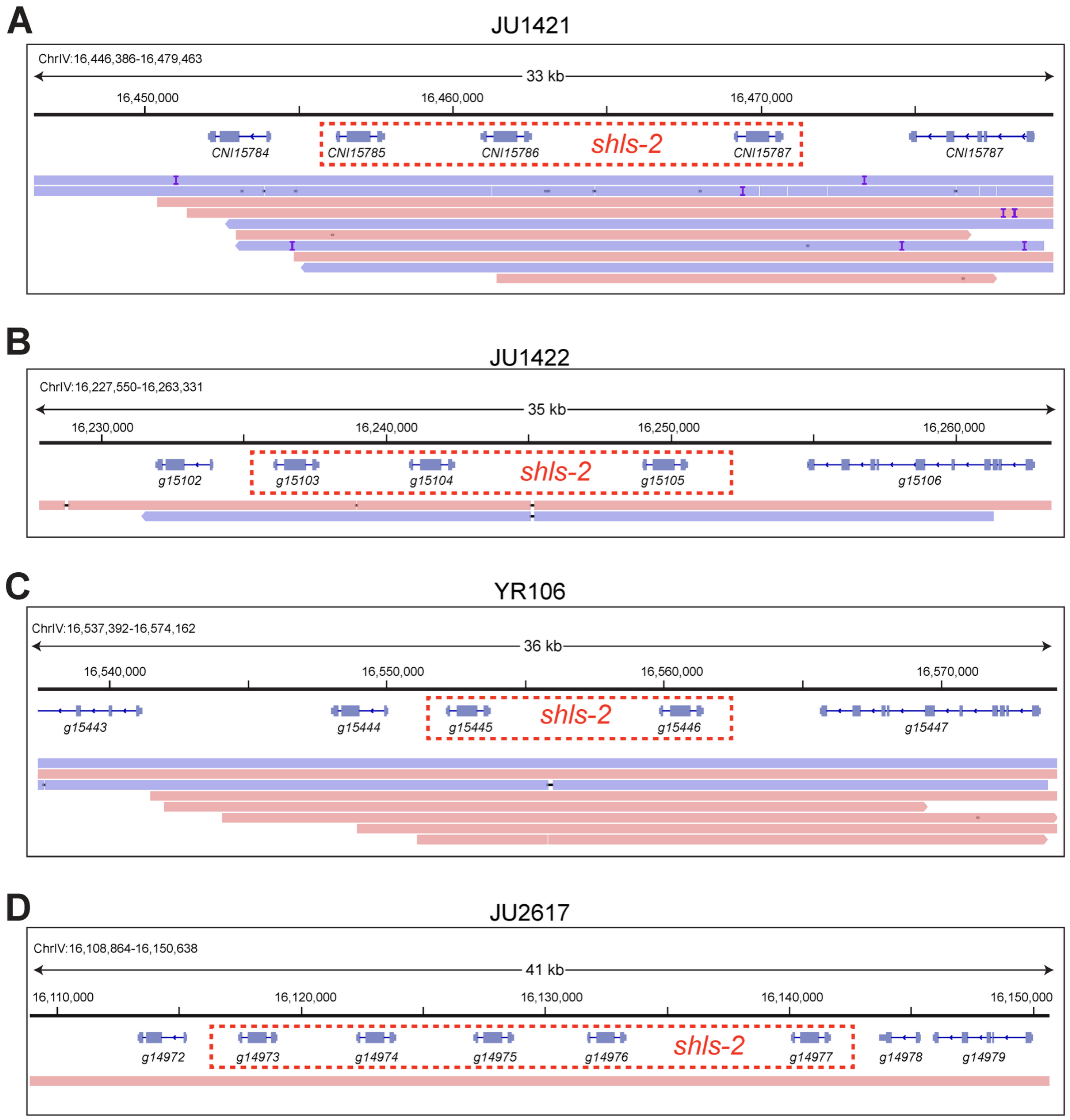


**Figure S14. ONT long-read sequencing confirms the duplication and copy number of *Cni-shls-2* in *C. nigoni* strains.**

**(A-D)** Genomic positions and predicted gene models of *Cni-shls-2* and adjacent genes are shown for strains JU1421 (A), JU1422 (B), YR106 (C), and JU2617 (D). Only ONT long reads spanning the entire *Cni-shls-2* genetic locus are displayed. Mapped reads are colored according to the mapped strand. The copies of *Cni-shls-2* paralogs are highlighted with red dashed rectangles.

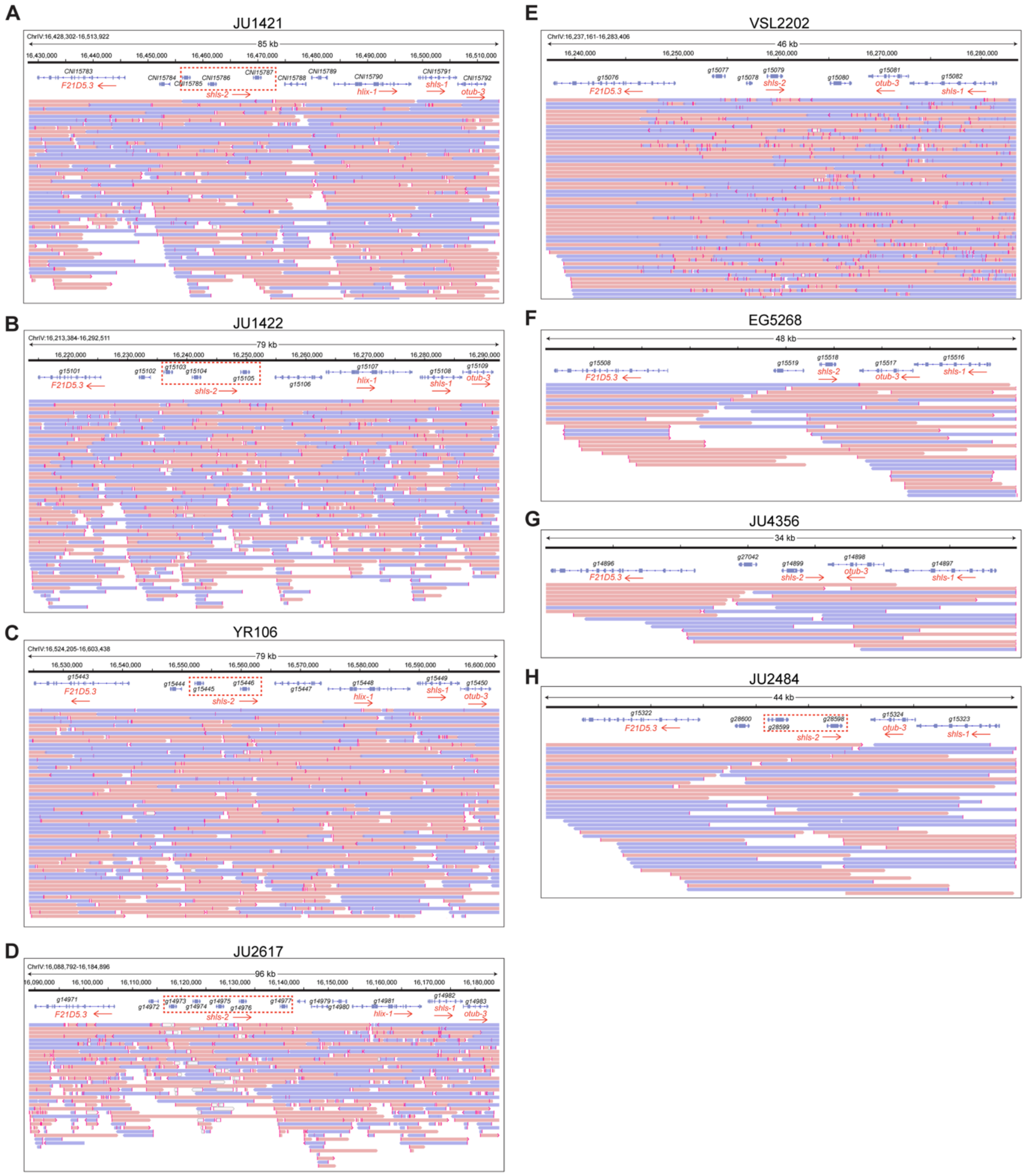


**Figure S15. ONT long-read sequencing confirms the relative positioning and orientation of the TA gene pair and its adjacent conserved genes, including *F21D5.3*, *shls-1*, and *otub-3*, in *C. nigoni* strains.**

**(A-H)** Genomic positions and predicted gene models for *Cni-shls-2* and adjacent genes are shown for strains JU1421 (A), JU1422 (B), YR106 (C), JU2617 (D), VSL2202 (E), EG5268 (F), JU4356 (G), and JU2484 (H). Mapped ONT long reads are displayed and differentially color-coded according to their mapped strand. *F21D5.3*, *Cni-shls-2*, *Cni-hlix-1* (if present), *Cni-shls-1*, and *otub-3* are labeled with their corresponding orientations, highlighted with red arrowheads. Note that the genomic regions for EG5268, JU4356, and JU2484 have been manually curated and are supported by ONT reads (see Methods).

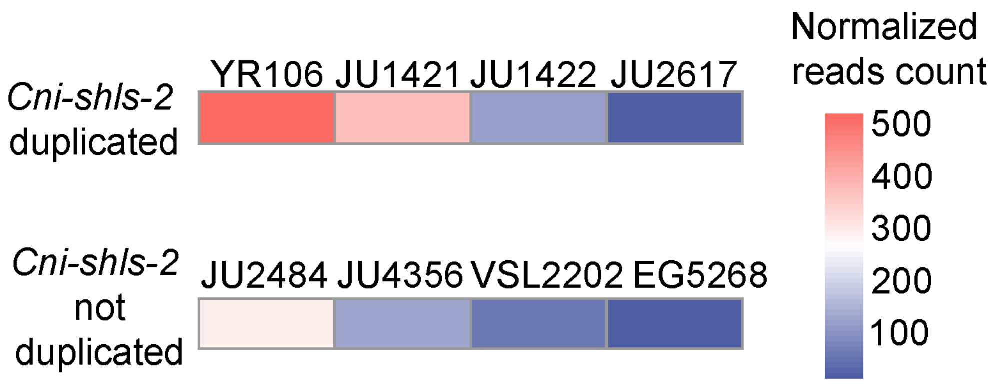


**Figure S16. Heatmap comparing the normalized expression levels of *Cni-shls-2* in *C. nigoni* strains carrying identical duplicated copies (top) versus those with a single copy (bottom).**

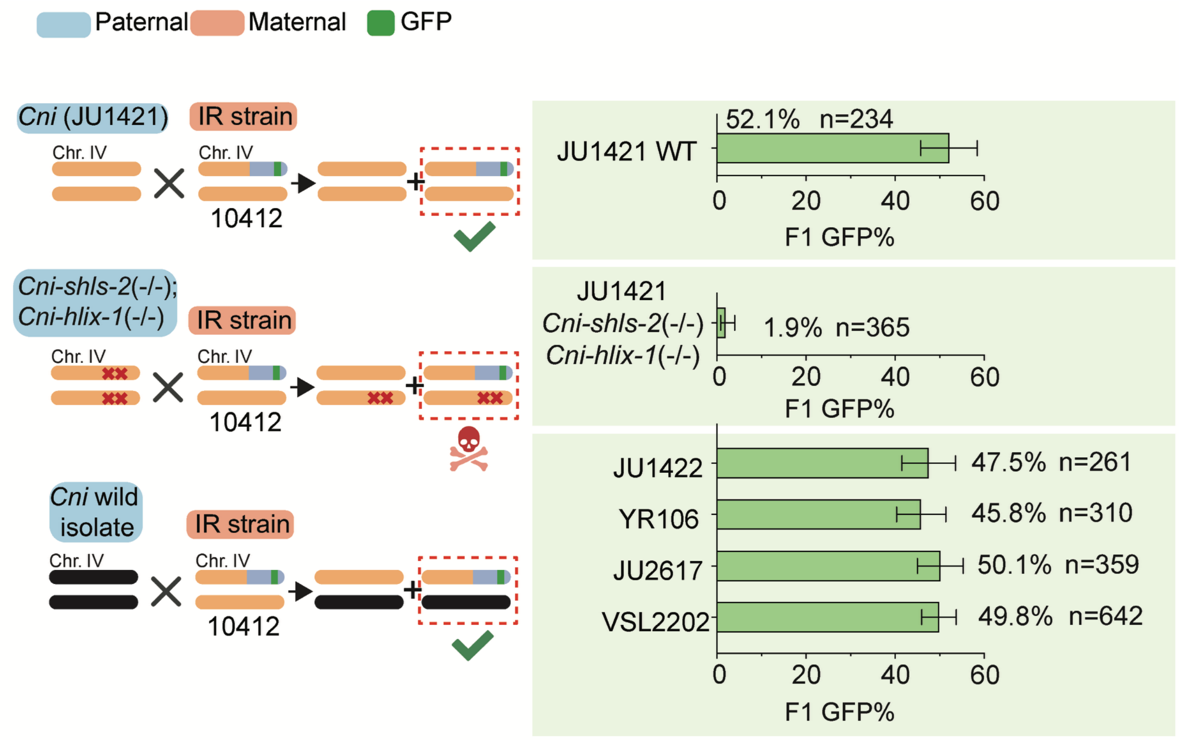


**Figure S17. *Cni-shls-2* from *C. nigoni* strains carrying the TA gene pair is functionally equivalent to that from JU1421.**

Comparison of the percentage of GFP-expressing adult F1 progeny from crosses between a wild-type *C. nigoni* (JU1421) father (top), a double homozygous TA gene mutant father in JU1421 (middle), or wild isolate *C. nigoni* fathers that carry a *Cni-shls-2* sequence 100%

identical to that of JU1421 *Cni-shls-2* (JU1422, YR106, JU2617 and VSL2202) (bottom), with a ZZY10412 mother. Error bars: 95% confidence intervals.

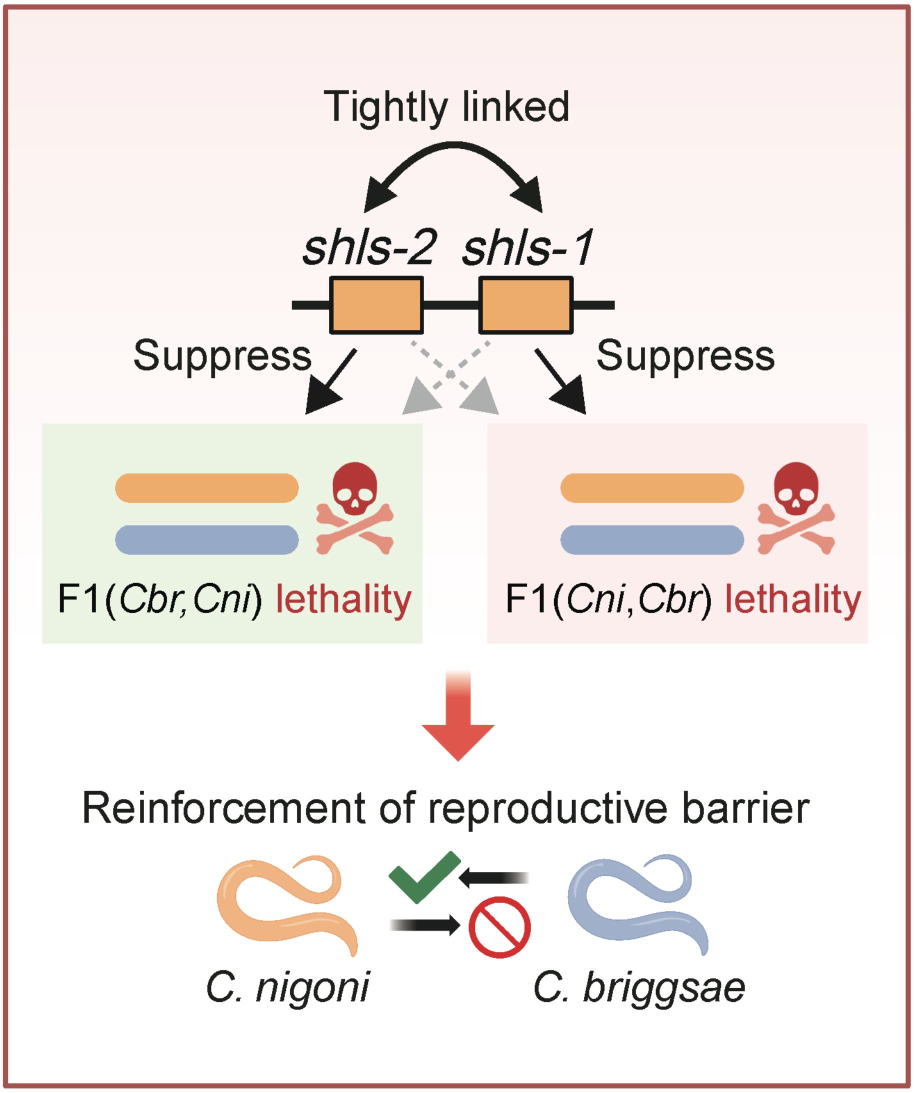


**Figure S18. Schematics illustrating how two tightly linked suppressor genes, each responsible for a distinct hybrid lethality mechanism, reinforce the reproductive barriers between *C. briggsae* and *C. nigoni*.**

*Cni-shls-1* suppresses hybrid lethality in progeny from a *C. briggsae* mother crossed with a *C. nigoni* father, but not in the reciprocal cross. In contrast, *Cni-shls-2* is responsible for hybrid lethality in progeny from a *C. briggsae* father and a *C. nigoni* mother, but not vice versa. The coexistence and tight linkage of these two suppressor alleles effectively block gene flow from *C. nigoni* to *C. briggsae*, thereby facilitating an asymmetric barrier that permits gene flow solely from *C. briggsae* to *C. nigoni* under laboratory conditions.
